# Hippo pathway and Bonus control developmental cell fate decisions in the *Drosophila* eye

**DOI:** 10.1101/2021.09.05.458934

**Authors:** Heya Zhao, Kenneth H. Moberg, Alexey Veraksa

## Abstract

The canonical function of the Hippo signaling pathway is regulation of organ growth. How this pathway controls cell fate determination is less well understood. Here, we uncover a function of the Hippo pathway in developmental cell fate decisions in the *Drosophila* eye-antennal disc exerted through the interaction of Yorkie (Yki) with the transcriptional regulator Bonus (Bon), an ortholog of mammalian Transcriptional Intermediary Factor 1/tripartite motif (TIF1/TRIM) family proteins. Instead of controlling tissue growth, Yki and Bon promote epidermal and antennal fates at the expense of the eye fate. Proteomic, transcriptomic, and genetic analyses reveal that Yki and Bon control these cell fate decisions by recruiting transcriptional and post-transcriptional co-regulators, and by activating epidermal differentiation genes and repressing Notch target genes. Our work expands the range of functions and regulatory mechanisms under Hippo pathway control.

## Introduction

Organ growth and cell fate determination are critical processes in organism development. These processes are controlled and kept in balance by multiple signaling pathways whose dysregulation leads to developmental abnormalities and diseases. One such pathway is the evolutionarily conserved Hippo pathway, which is implicated in various physiological functions and pathological conditions including organ growth, cell differentiation, tissue regeneration, and oncogenesis (Kulkarni et al., 2020; Zheng and Pan, 2019). The Hippo pathway transduces upstream regulatory signals to transcriptional outputs that mediate various cellular functions including cell proliferation and survival (Meng et al., 2016). The core Hippo pathway acts via the Hippo (Hpo)/Warts (Wts) kinase cascade to negatively regulate the activity of the transcriptional coactivator Yorkie (Yki), which associates with DNA-binding proteins such as Scalloped (Sd) to control gene expression (Zheng and Pan, 2019). The canonical transcriptional targets of the Yki-Sd complex in *Drosophila* include *cyclin E* (*cycE*), *death-associated inhibitor of apoptosis 1* (*diap1*), *bantam* microRNA (*mir-ban*), and *expanded* (*ex*); these factors promote proliferation, inhibit apoptosis, and enable negative feedback regulation (Pan, 2010). Although increasing evidence supports an essential role of the Hippo pathway in cell fate determination in multiple biological contexts (Davis and Tapon, 2019), the cellular mechanisms remain poorly understood.

The *Drosophila* eye is an excellent model to study gene regulatory networks controlling cell fate determination (Weasner and Kumar, 2022). Most of the *Drosophila* adult head structures develop from the larval eye-antennal disc, with the compound eye and ocelli originating from the eye disc compartment, the antenna and maxillary palp from the antennal compartment, and the head epidermis from the tissues surrounding the two compartments (Dominguez and Casares, 2005). Segregation of the mutually antagonistic eye, antennal, and head epidermal fates, which begins at the second instar larval stage (L2), is regulated by several signaling pathways including Notch, EGFR, Wingless, and Hedgehog, and retinal determination genes such as *eyeless* (*ey*) and *homothorax* (*hth*) (Kumar, 2010; 2018). Alteration of these regulatory inputs can cause a switch from one fate to another, leading to partial or in some cases complete homeotic transformations of the affected structures (Kumar and Moses, 2001; Ordway et al., 2021; Wang and Sun, 2012; Weasner and Kumar, 2013). Key patterning events in the eye are linked to a wave of differentiation called the morphogenetic furrow (MF) that starts in the early third instar larval stage (L3) and proceeds from the posterior to the anterior of the eye disc, resulting in the differentiation of an array of optical units called ommatidia, each consisting of photoreceptor cells, cone cells, primary pigment cells, interommatidial bristles, and secondary and tertiary pigment cells (also called interommatidial cells) (Cagan and Ready, 1989a; Charlton-Perkins and Cook, 2010).

Previous studies of the Hippo pathway in *Drosophila* eye differentiation focused on MF progression, terminal differentiation of photoreceptor cells, and formation of peripodial epithelium (Jukam et al., 2013; Pojer et al., 2021; Wittkorn et al., 2015; Zhang et al., 2011). Mutant analyses of the Hippo pathway components *ex*, *Merlin* (*Mer*) and *mob as tumor suppressor* (*mats*) have suggested an earlier and broader impact of the Hippo pathway in eye specification (Boedigheimer and Laughon, 1993; Lai et al., 2005; McCartney et al., 2000; Pellock et al., 2007). However, the involvement of the Hippo pathway in controlling cell fate decisions among the eye, antenna, and head epidermis remains elusive, and the underlying transcriptional mechanisms are unknown.

We reasoned that the Hippo pathway may have a novel function in controlling the eye-antenna-epidermis fate determination through yet unknown interactors that regulate the transcriptional output of the Yki-Sd complex. To identify such interactors, we performed proteomic analyses and identified a novel Yki-interacting protein, Bonus (Bon). Bon is the only *Drosophila* ortholog of mammalian TIF1 family proteins TIF1α (TRIM24), TIF1β (TRIM28/KAP1), TIF1γ (TRIM33), and TIF1δ (TRIM66) (Beckstead et al., 2001; Cammas et al., 2012). TIF1/Bon proteins are chromatin-associated factors that activate or repress transcription by binding to co-regulators and controlling chromatin state (Beckstead *et al*., 2001; Beckstead et al., 2005; Cammas *et al*., 2012; Ito et al., 2012). TIF1 proteins play various roles during vertebrate development and are implicated in cancer (Cambiaghi et al., 2012; Petrera and Meroni, 2012). Invertebrate Bon is essential for nervous system development, metamorphosis, and cell survival (Allton et al., 2009; Beckstead *et al*., 2001; Ito *et al*., 2012; Salzberg et al., 1997).

Here, we present evidence that Bon and the Hippo pathway co-regulate major cell fate decisions during the development of the *Drosophila* eye. Yki and Bon bind via WW domain-PPxY motif interactions and cooperate to produce epidermal cells in the eye at the expense of ommatidial cells, while loss of *bon* induces ectopic eye markers, suggesting that the Hippo pathway and Bon control the choice between the eye and epidermal fates. The Hippo pathway and Bon also regulate the eye-antennal specification, with Yki and Bon inhibiting the eye fate and promoting the antennal fate. Through the analysis of Bon and Yki protein interactors, we have identified multiple transcriptional and post-transcriptional regulators that are necessary for their control of cell fate decisions. Transcriptome analysis has revealed that Bon and Yki exert their functions by jointly activating epidermal differentiation genes and, unexpectedly, repressing Notch target genes. Overall, we have identified a novel function of the Hippo pathway in the eye/antenna/epidermis cell fate decisions during *Drosophila* eye development. This function requires the interaction of Yki with Bon, their recruitment of coregulators, and the joint transcriptional control of a non-canonical set of target genes.

## Results

### Bon is a novel Yki interactor

To identify new regulators of the Hippo pathway, we performed affinity purification-mass spectrometry (AP-MS) using *Drosophila* embryos expressing Yki-EGFP with a ubiquitous driver *da-GAL4* (Wodarz et al., 1995), as well as cultured *Drosophila* S2 cells expressing streptavidin-binding peptide (SBP)-tagged Yki (see Method Details). We identified the core Hippo pathway components, accessory regulators, and several putative novel Yki interactors, including Bon (Figure 1A, Table S1) (Kwon et al., 2013; Oh et al., 2014; Zheng and Pan, 2019). We confirmed the interaction between Bon and Yki by co-immunoprecipitation (co-IP) of tagged Bon and Yki proteins in S2 cells (Figures 1C-D). We hypothesized that the Yki-Bon interaction may be mediated by the two PPxY motifs in Bon (PPLY^507^ and PPSY^585^) binding to the WW domains in Yki (Figure 1B), similar to other known Yki interactors (Huang et al., 2005; Oh *et al*., 2014). Mutating the key tyrosine residue to alanine (Zhang et al., 2015) in either single or both WW domains in Yki strongly reduced its binding to Bon, with the first WW domain (WW1) showing a stronger reduction in binding upon mutation (Figure 1C). Similarly, substituting tyrosine with alanine in one or both PPxY motifs in Bon also largely reduced its binding to Yki, with a stronger reduction by the mutation in the second PPxY motif (PPxY2) (Figure 1D). These results indicate that Bon interacts with Yki through PPxY motifs in Bon and WW domains in Yki.

**Figure 1.**
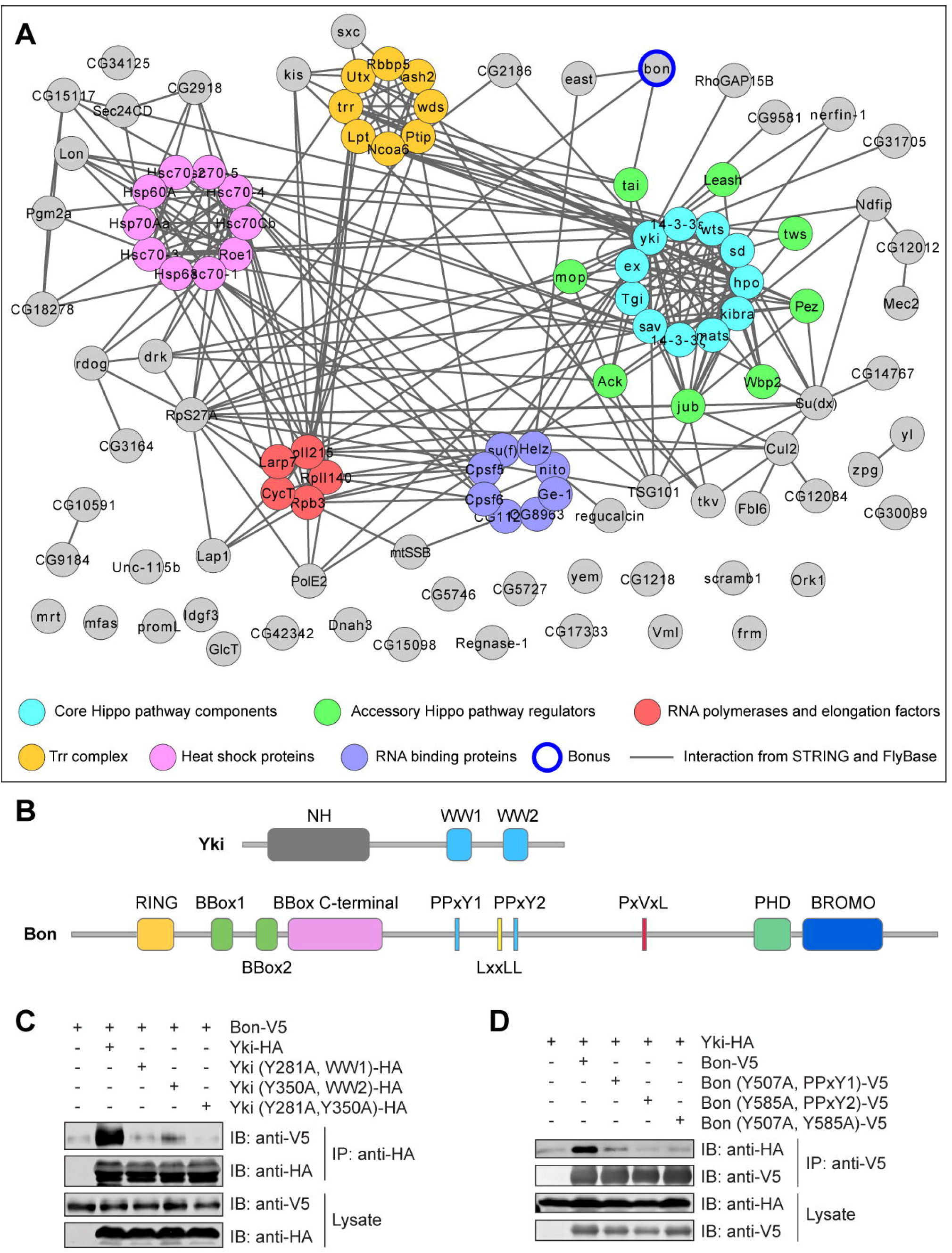
Identification of Bonus as a Yorkie interactor. (A) The Yki protein interaction network. AP-MS results of Yki-SBP from *Drosophila* S2 cells and Yki-EGFP from embryos were analyzed with Significance Analysis of INTeractome (SAINT) (Choi et al., 2011). All the interactors (nodes) in this network were identified as significant Yki interactors (SAINT score ≥ 0.8); these interactions (edges) are not shown to avoid clutter. The shown edges were incorporated from the STRING database and FlyBase, and include various types of evidence (Szklarczyk et al., 2019; Thurmond et al., 2019). Also see Method Details. (B) Schematic diagram showing the major domains and motifs in Yki and Bon. Note two WW domains in Yki and two PPxY motifs in Bon. (C) Co-IP of Bon-V5 and wild-type or mutant Yki-HA expressed in S2 cells. (D) Co-IP of Yki-HA and wild-type or mutant Bon-V5 expressed in S2 cells.

### Activation of Bon or Yki induces epidermal trichomes in adult eyes

We next sought to investigate whether Bon is involved in Yki-mediated growth regulation. RNAi knockdown of *bon* did not affect the overgrowth or cell proliferation induced by Yki overexpression in adult eyes, larval eye discs, and wing discs (Figures S1A-D). In addition, RNAi knockdown of *bon* in wing discs did not affect the reporters of canonical Yki target genes *diap1* and *ex,* which mediate Yki’s function in growth regulation through anti-apoptosis and feedback regulation (Figure S1E). Loss-of-function *bon* mutants exhibit ectopic abdominal peripheral neurons in the embryo (Beckstead *et al*., 2001; Salzberg *et al*., 1997). Null mutants of *wts* (*wts^X1^*) (Xu et al., 1995) and *yki* (*yki^B5^*) (Huang *et al*., 2005) did not show loss or gain of peripheral neurons and did not affect the ectopic neuron phenotype in null (*bon^21B^*) or hypomorphic (*bon^487^*) *bon* mutant embryos (Figure S1F). Collectively, these data argue that Yki and Bon have independent functions in growth regulation and embryonic peripheral nervous system (PNS) development.

Interestingly, overexpression of Bon with *GMR-GAL4*, which drives expression in all cells after the MF and persists through pupal eye development (Ellis et al., 1993; Hay et al., 1994), resulted in the formation of trichomes on the surface of adult eyes and disruption of the ommatidial array (Figures 2A-B and S2A), which is consistent with a previous report (Tardi et al., 2012). Trichomes are actin-rich, non-neuronal apical extensions, which are present on *Drosophila* epidermal cells but not retinal cells (Arif et al., 2015; Delon et al., 2003). This trichome induction suggests that Bon promotes epidermal fate in the eye.

**Figure 2.**
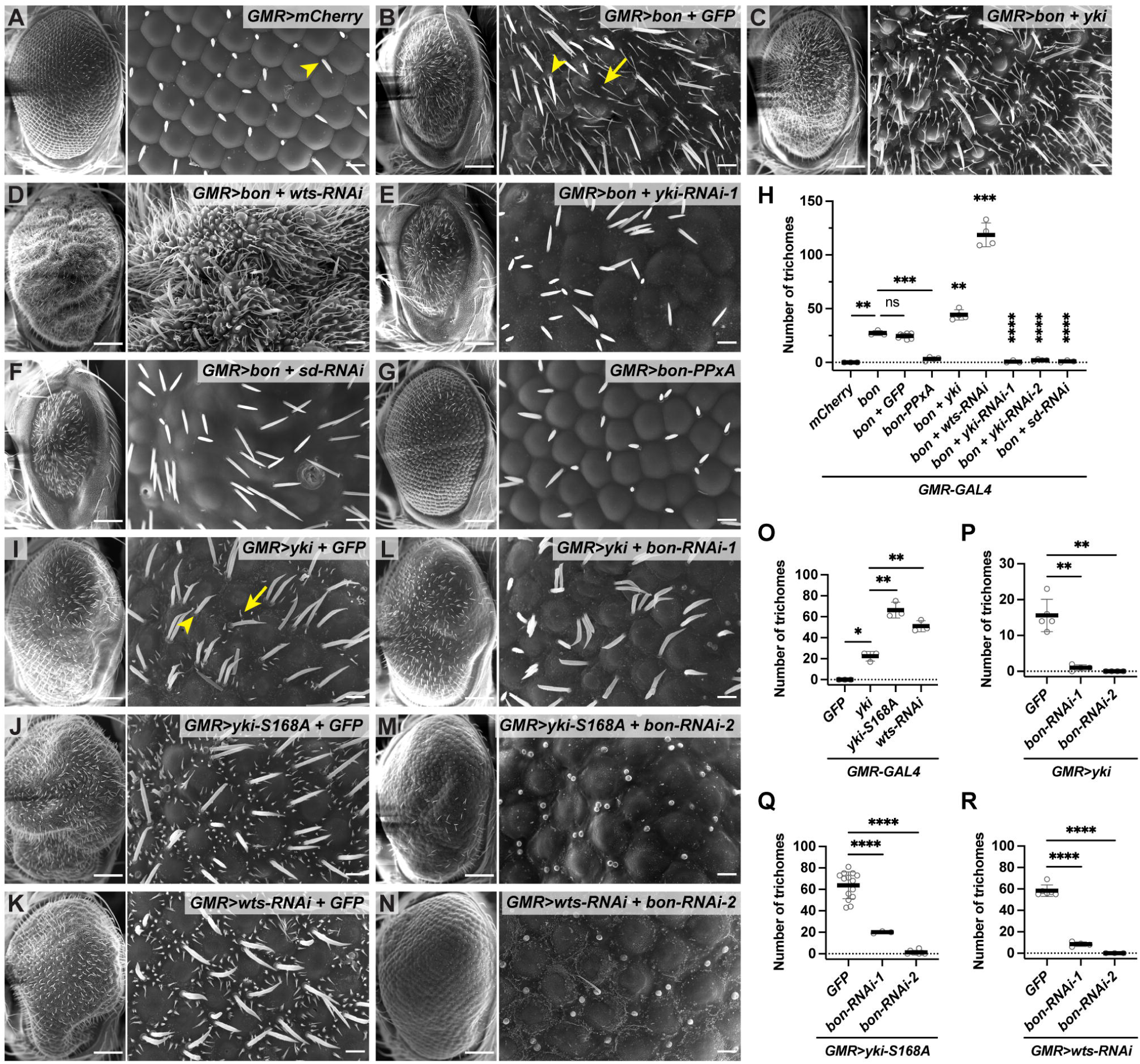
Activation of Bon or Yki induces epidermal trichomes in adult eyes. (A-G, I-N) SEM images of adult *Drosophila* eyes expressing indicated *UAS* transgenes with the *GMR-GAL4* driver. Crosses with Bon were carried out at 25°C, and crosses with Yki, Yki-S168A and *wts-RNAi* were set up at 25°C and shifted to 29°C after the emergence of first instar larvae. Arrowheads: interommatidial bristles, arrows: trichomes. Scale bars in left panels: 100 μm; enlarged views in right panels: 10 μm. (H, O-R) Quantification of trichome numbers in A-G (H) and I-N (O-R). *GFP* was coexpressed for transgene dosage compensation. Trichome numbers here and in the other figures were obtained in an area of 1306 μm^2^. Two additional *bon-RNAis* were used that significantly reduced trichome numbers, with images and quantifications given in Figure S2 and Table S5, respectively. Images for other quantified genotypes are shown in Figure S2. The number of trichomes and the number of images quantified (n ≥ 3) for each genotype are provided in Table S5. Statistical analyses are described in Method Details.

To examine whether the Hippo pathway is involved in Bon trichome induction, we overexpressed or knocked down key components of the Hippo pathway in the Bon overexpression background and quantified trichome number. Yki overexpression or *wts* knockdown increased Bon-induced trichomes (Figures 2C-D and 2H). In contrast, knockdown of *yki* or *sd* strongly suppressed Bon-induced trichomes, with only mild eye roughness observed in single knockdowns (Figures 2E-F, 2H, S2B, and S2R-T). Thus, the core Hippo pathway and Yki transcriptional activity are essential for Bon-induced trichomes in adult eyes. Furthermore, overexpression of Bon-PPxA, a Bon mutant with both PPxY motifs mutated to PPxA that cannot bind to Yki (see Figure 1D), resulted in significantly fewer trichomes and milder ommatidial defects (Figures 2G-H), indicating that Bon must bind Yki to efficiently induce eye trichomes.

We then investigated whether Yki activation is sufficient to induce trichomes in adult eyes. Indeed, Yki overexpression by *GMR-GAL4* resulted in trichome formation in the eye (Figures 2I and S2C-D). Overexpression of constitutively active Yki-S168A, a Yki mutant insensitive to Wts phosphorylation and inhibition (Oh and Irvine, 2008), induced even more trichomes (Figures 2J, 2O, and S2G). Knockdown of *wts* also induced more trichomes than Yki overexpression (Figures 2K, 2O, and S2K). Therefore, Yki activation promotes trichome formation, with trichome number positively correlated with Yki activity. Knockdown of *bon* strongly suppressed trichome formation induced by wild-type Yki, Yki-S168A, and *wts*-RNAi, with only mild roughness when knocked down alone (Figures 2L-N, 2P-R, and S2D-Q), suggesting that Bon is necessary for trichome formation induced by activated Yki. Together, these results show that the Hippo pathway and Bon jointly regulate the epidermal trichome formation in the eye.

### Hippo pathway and Bon control the decision between eye and epidermal cell fates

Next, we investigated cell fate changes underlying ectopic trichome induction in the retina. The establishment of trichomes occurs at 30-36 hrs after puparium formation (APF) in the wing and 38-39 hrs APF in the notum (Chanut-Delalande et al., 2014; Delon *et al*., 2003). F-actin and cell boundary (Dlg) staining of pupal eyes revealed that Bon-induced trichomes are initiated at 40 hrs APF (Figure S3A-C) and are easily visible at 44 hrs APF (Figures 3A). Bon overexpression resulted in severe disruption of the ommatidia and produced extra interommatidial-like cells from which the trichomes derive (Figure 3A). Importantly, the excess interommatidial-like cells were generated without alteration in DNA synthesis (Figure S1D) or overall apical cell numbers (Figure S3E), suggesting a failure of differentiation rather than aberrant proliferation or survival. Interommatidial cells are believed to be the default state in retinal differentiation (Charlton-Perkins and Cook, 2010; Kumar, 2012). Trichomes on these cells indicate that they may be reprogrammed into epidermal-like cells. Knockdown of *yki* or *sd* strongly suppressed the initiation of Bon-induced trichomes and partially restored the ommatidia (Figure 3A). Therefore, Bon induces epidermal cell fate at the expense of the eye fate in a Yki and Sd-dependent manner.

**Figure 3.**
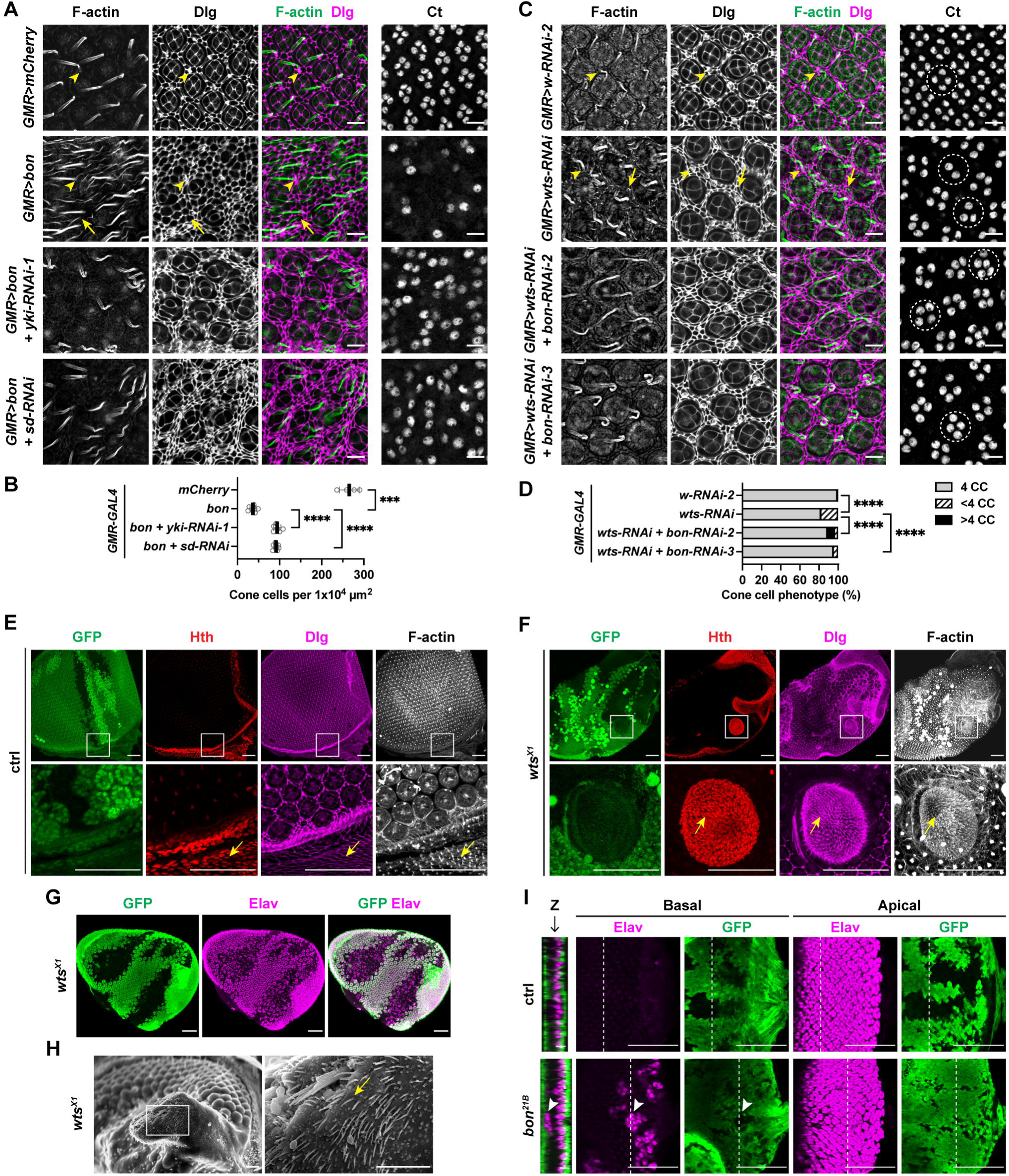
Yki and Bon promote the epidermal fate at the expense of the eye fate. (A, C) Pupal eyes expressing the indicated transgenes with *GMR-GAL4* were stained with phalloidin for F-actin and anti-disc large (Dlg) antibody for cell boundaries, or anti-Cut (Ct) antibody for cone cells. (A) 44 hrs APF grown at 25°C, (C) 40 hrs APF grown at 29°C (equivalent to 48 hrs APF at 25°C). Arrowheads: interommatidial bristles and sockets, arrows: trichomes and the corresponding cells, dashed circles in C: individual cone cell clusters per ommatidium. Scale bars: 10 μm. (B, D) Quantification of cone cell numbers per area in (A) or per ommatidium in (C), respectively. Cone cell counts in (B) were done in an area of 1×10^4^ μm^2^ instead of per ommatidium due to a severe phenotype. (D) The *p* values were determined using Fisher’s exact test between ommatidia with ≥4 cone cells (CC) and those with <4 CC per ommatidium. The numbers are provided in Table S5. (E-F) Pupal eyes at 46 hrs APF with mosaic clones of the indicated genotypes were immunostained with anti-Hth antibody for head epidermis, anti-Dlg antibody for cell boundaries, and phalloidin for F-actin. Mutant clones were generated with heat shock at 39 hrs after egg deposition (AED) and marked by loss of GFP. Enlarged views of the boxed regions are shown in the bottom panels. Arrows: trichomes and the corresponding cells in wild type (WT) head cuticle (E) or ectopic epidermis in the eye (*wts^X1^* clone in F). Scale bars: 50 μm. Exact genotypes of mutants and reporters here and in other figures are given in Method Details. (G) Pupal eye at 46 hrs APF with mosaic clones of *wts^X1^*immunostained with anti-GFP antibody to mark the cells outside the clones and anti-Elav antibody for neuronal eye fate. Mutant clones were generated with heat shock at 42 hrs AED. Scale bars: 50 μm. (H) Representative SEM image of adult eye with mosaic clones of *wts^X1^*. Mutant clones were generated with heat shock at 39 hrs AED. Enlarged view of the boxed region is shown in the right panel. Arrow: trichome. Scale bars: 20 μm. (I) L3 eye disc with mosaic clones of the indicated genotypes immunostained with anti-GFP and anti-Elav antibodies. Mutant clones were generated with *eyFLP*. Orthogonal sections (Z) at the dashed lines of basal/apical (2D) views are shown in the left panels. Arrows: the ectopic eye (Elav) and the corresponding clone (loss of GFP) in the basal layer of the *bon^21B^* clone. The orthogonal sections and their scale bars were scaled 2x along the z axis for easier visualization. Scale bars: 5 μm in Z views and 50 μm in 2D views.

To determine more precisely which retinal cells were affected, we immunostained pupal eyes at 44 hrs APF for cone cell marker Ct, primary pigment cell marker BarH1, and photoreceptor cell and bristle group marker Elav (Charlton-Perkins and Cook, 2010). Bon overexpression with *GMR-GAL4* strongly inhibited formation of cone cells and primary pigment cells, and affected patterning of photoreceptor cells and bristle groups (Figures 3A-B and S3D). *yki* or *sd* knockdown in this background suppressed the loss of cone cells and partially restored the mispatterned photoreceptor cells and bristle groups (Figures 3A-B and S3D), but did not rescue the loss of primary pigment cells (Figure S3D), likely because the differentiation of primary pigment cells requires successful differentiation of cone cells (Nagaraj and Banerjee, 2007). These results demonstrate that Bon, Yki, and Sd jointly suppress eye fate.

*wts* knockdown by RNAi also induced trichomes on interommatidial cells at mid-pupal stage, suggesting that these cells were reprogrammed into the epidermal fate (Figure 3C). Although the trichomes induced by *wts* RNAi were shorter and thicker than Bon-induced trichomes (Figures 2B and 2K), they were clearly distinguishable from the bristle shafts of the bristle groups which have sockets (Figure 3C, Dlg staining) and neuronal input (Figure S3I, Elav staining) (Charlton-Perkins and Cook, 2010). *bon* knockdown strongly suppressed *wts* RNAi-induced trichome initiation, but not the excess interommatidial cells and bristles which result from overproliferation (Huang *et al*., 2005; Udan et al., 2003), suggesting that Bon is specifically necessary for cell fate reprogramming (Figure 3C). Consistent with previous studies, *wts* RNAi reduced the number of cone cells (Hamaratoglu et al., 2006; Wittkorn *et al*., 2015), which was rescued by knockdown of *bon* (Figures 3C-D). Moreover, *bon* knockdown alone resulted in gain of cone cells, especially at the periphery of the retina (Figures S3F-G). Loss of *wts* and *bon* had a mild effect on photoreceptor patterning and showed a mixed outcome for primary pigment cells (either gain or loss per ommatidium) (Figure S3H). Together, these results show that Bon activation or Hippo pathway inactivation behind the MF induces epidermal fate at the expense of the eye fate, as represented by trichome formation and inhibition of cone cells.

We further investigated endogenous control of cell fate determination using loss of function clones of *wts* and *bon* alleles. Trichomes were apparent in *wts^X1^* eye clones at 46 hrs APF, and the trichome-making cells strongly expressed Hth, which normally marks the head epidermis at this stage (Figure 3E-F) (Lim and Tomlinson, 2006; Wernet et al., 2003). *wts^X1^* clones also suppressed eye fate, as evidenced by loss of ommatidia (Dlg in Figure 3F) and photoreceptors (Elav in Figure 3G). In adult eyes, *wts^X1^* clones produced outgrowths of epidermal tissue covered with trichomes (Figure 3H). Interestingly, *bon^21B^* clones led to gain of neuronal eye marker Elav at the basal side of the larval eye disc, without affecting the normal apical patterns of Elav, suggesting an induction of ectopic eye tissue (Figure 3I). Altogether, our results using both loss and gain of function analyses suggest that Bon and Yki (or inactivated core Hippo pathway) suppress eye fate and promote the epidermal fate.

### Hippo pathway and Bon regulate the choice between eye and antennal cell fates

Since eye fate is antagonistic to epidermal and antennal fates (Weasner and Kumar, 2022), we asked whether Hippo and Bon also regulate the balance of eye-antennal fates. To explore this hypothesis, we used the early eye driver *ey-GAL4* which is expressed in the entire eye disc beginning at the L2 stage, and examined L3 eye-antennal discs using Hth and Ct as antennal markers, and Elav as the differentiated neuron/eye marker (Blochlinger et al., 1993; Kumar and Moses, 2001; Pai et al., 1998; Wang and Sun, 2012). *wts* knockdown or Yki-S168A overexpression by *ey-GAL4* significantly suppressed the eye field, with the respective penetrance of 56% (*wts*-RNAi), 100% (*yki-S168A*-1), and 85% (*yki-S168A*-2), and even occasionally led to a complete eye-to-antenna transformation, with the respective penetrance of 11%, 5%, and 6% (Figures 4A-C, 4I, S4A-C, S4E, S4I-K and Table S5). Remarkably, *wts* knockdown or Yki-S168A overexpression also led to clear eye-to-antenna transformations in some adult flies (Figures 4E-G). Wildtype Yki overexpression with *ey-GAL4* suppressed the eye field and resulted in extra bristles (vibrissae) and occasional epidermal outgrowths, but without eye-to-antenna transformation (Figures S4G-G’’’).

**Figure 4.**
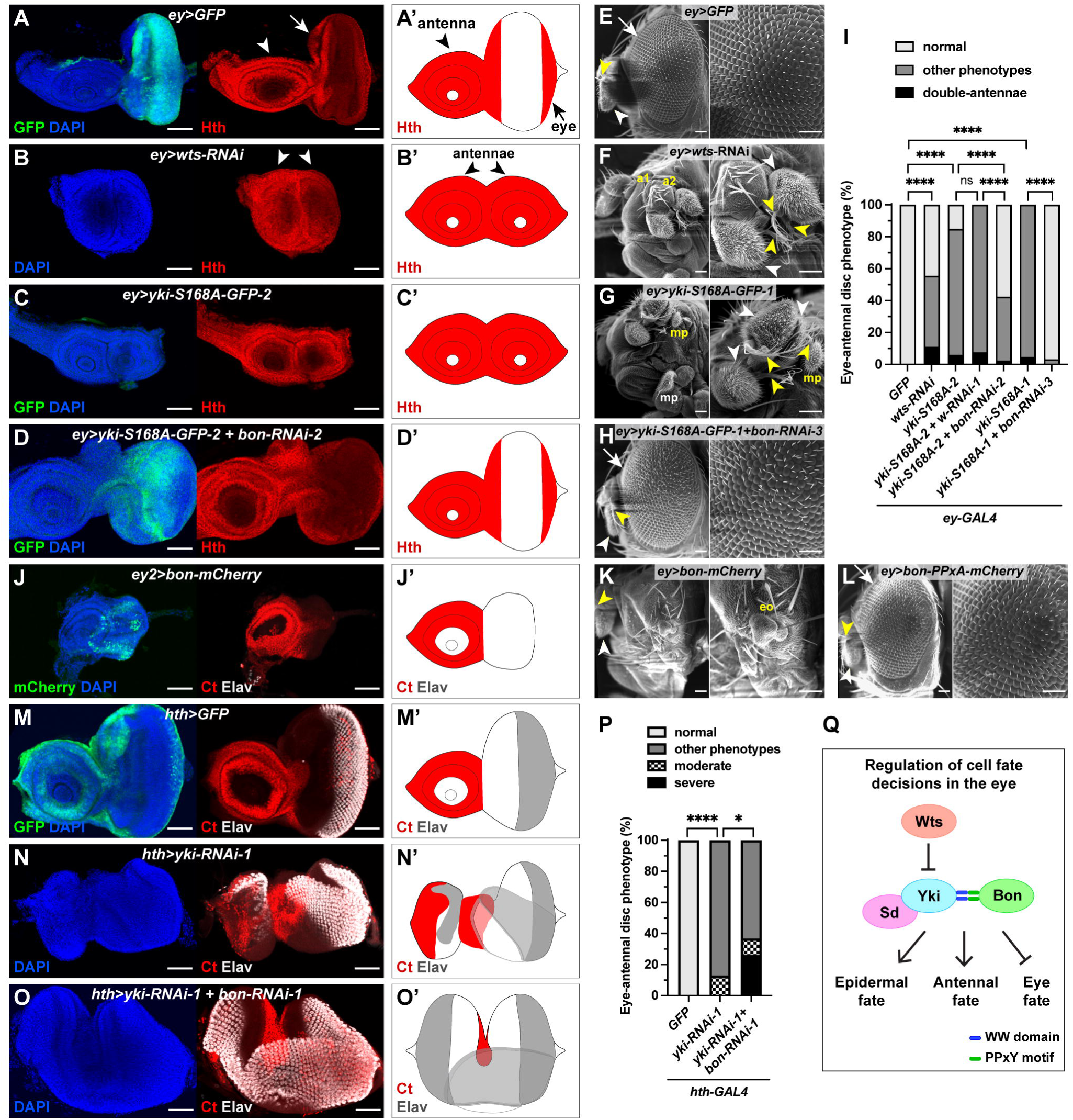
Yki and Bon promote antennal fate and suppress eye fate. (A-D’) L3 eye-antennal discs expressing indicated transgenes with *ey-GAL4* were immunostained with anti-Hth antibody for the antennal field. Expression region of the driver is visualized by GFP (A). (A’-D’) Schematic illustrations for the structure of the eye-antennal disc and the pattern of Hth staining in A-D. Arrows: eye discs, arrowheads: antennal discs. Scale bars: 50 μm. (E-H) Representative SEM images of adult heads expressing the indicated transgenes with *ey-GAL4*. (F) *wts-RNAi* had double antennae with triple aristae and fused a1 and a2 segments. (G) *Yki-S168A* had double or triple antennae with one arista each and an ectopic maxillary palp. Right panels: enlarged views of the phenotypes, arrows: eyes, white arrowheads: antennal a3 segments, yellow arrowheads: aristae, white mp and yellow mp: endogenous and ectopic maxillary palps, respectively. Scale bars: 50 μm. (I) Quantification of the phenotypes in eye-antennal discs shown in A-D, S4A-S4F and S4I-S4K. P values were determined using Fisher’s exact test between normal and abnormal eye disc populations within indicated comparisons. The numbers are provided in Table S5. (J-J’) L3 eye-antennal disc expressing *bon-mCherry* with *ey-GAL4-2* immunostained with anti-Ct antibody for the antennal field and anti-Elav antibody for the neuronal eye fate. mCherry and Ct signals were colored green and red, respectively. (J’) Schematic illustration of J. Control with GFP expressed under the same driver is shown in S4O-O’. Quantifications are provided in Table S5. Scale bars: 50 μm. (K-L) Representative SEM images of adult heads expressing the indicated transgenes with *ey-GAL4*. Right panels: enlarged views of the phenotypes, arrow: eye, white arrowhead: antennal a3 segment, yellow arrowhead: arista, eo: epidermal outgrowth. Quantifications are provided in Table S5. Scale bars: 50 μm. (M-O’) L3 eye-antennal discs expressing indicated transgenes with *hth-GAL4* were immunostained with anti-Ct and anti-Elav antibodies. Expression region of the driver in disc proper is shown with GFP (M). Representative moderate (N) and severe (O) antenna-to-eye transformations are shown. (M’-O’) Schematic illustrations of M-O. Scale bars: 50 μm. (P) Quantification of the phenotypes in eye-antennal discs shown in M-O and S4L-S4N. P values were determined using Fisher’s exact test between normal and abnormal antennal disc populations (*GFP* vs. *yki-RNAi-1*) or between severe transformation and non-severe phenotypes (*yki-RNAi-1 + bon-RNAi-1* vs*. yki-RNAi-1*). The numbers are provided in Table S5. (Q) Summary model showing the function of Wts, Yki, Sd and Bon during cell fate decisions in the eye: the Yki-Bon complex promotes epidermal fate and antennal fate, and suppresses eye fate.

*bon* knockdown significantly rescued the smaller eye field resulting from Yki-S168A overexpression (from 100% to 3%, and 85% to 43%, with two independent lines) and suppressed formation of double antennae (from 5% to 0%, and 6% to 2.5%, with two lines) (Figures 4D, 4I, S4D, S4F, and Table S5). In adults, *bon* knockdown also largely rescued Yki-S168A-induced loss of the eye (Figure 4H). Bon overexpression using *ey-GAL4* suppressed the differentiated eye field in eye-antennal discs with a penetrance of 20% (Figures 4J, S4O, and Table S5), and led to a complete or partial loss of the eye with frequent epidermal outgrowths in adults, with a penetrance of 18% (Figure 4K and Table S5), but without eye-to-antenna transformation. In contrast, Bon-PPxA overexpression led to largely normal eyes (Figure 4L and Table S5). Altogether, these results show that early and strong activation of Yki across the eye field can switch the eye fate to antennal fate in a Bon-dependent manner, while weaker activation of Yki or Bon transforms eye to epidermis.

To determine whether the eye vs. antennal fate choice is under control of endogenous *yki* and *bon*, we tested effects of *yki/bon* loss in the antenna using the *hth-GAL4* driver (Dominguez and Casares, 2005; Wernet *et al*., 2003). Since eye and antennal fates are mutually antagonistic and reciprocally transformable (Wang and Sun, 2012), we hypothesized that loss of *yki* or *bon* in antenna would promote eye fate. Indeed, *yki* knockdown alone resulted in variable degrees of antenna loss with 100% penetrance, and moderate antenna-to-eye transformations (13% penetrance) (Figures 4M-N, 4P, S4L-N, and Table S5). This partial antenna-to-eye transformation exhibited loss of Ct and gain of Elav in the antennal field, suggesting a suppression of the antennal fate and differentiation of ectopic eye tissue (Figure 4N). Strikingly, a double knockdown of *yki* and *bon* resulted in a severe antenna-to-eye transformation with a 26% penetrance, in which the antennal field marked by Ct was restricted to the center of the eye-antennal disc, and the ectopic Elav pattern resembled a mirror-image duplication of the endogenous eye field (Figures 4O-P). These results demonstrate that endogenous Yki and Bon are necessary for the specification of the antennal fate and the suppression of the eye fate. Altogether, our results support a model of eye specification regulated by the Hippo pathway and Bon, where Yki and Bon promote epidermal fate and the antennal fate, while suppressing eye fate (Figure 4Q).

### Interactors of Bon and Yki are required for the eye-epidermal fate choice

We sought to further investigate the mechanism through which Yki and Bon regulate the eye-epidermal fate choice. One potential mechanism is that Bon enhances Yki activity by competing Yki away from Wts, given that both Bon and Wts bind to Yki through the PPxY-WW interaction. However, co-IP in S2 cells showed no competition between Bon and Wts for binding to Yki (Figure S5A). This result suggests that rather than competing with Wts, Bon may employ another mechanism, such as engaging a special set of interactors to mediate the eye-epidermal cell fate decision together with Yki.

To identify Bon cofactors, we analyzed the Bon interactome by AP-MS using Bon-SBP expressed in S2 cells (Figure 5A and Table S2). Yki was identified as one of the top Bon interactors, further validating their interaction (Figure 5A). In addition, we identified Histone deacetylase 1 (HDAC1), Suppressor of variegation 2-10 (Su(var)2-10), and Heterogeneous nuclear ribonucleoprotein at 27C (Hrb27C) as top Bon interactors (Figure 5A). HDAC1 is a transcriptional corepressor that is a key component of several chromatin remodeling complexes, such as NuRD, Sin3, and CoRest (Seto and Yoshida, 2014). Su(var)2-10 belongs to the PIAS family of SUMO ligases and controls gene transcription as well as *Drosophila* eye specification (Betz et al., 2001; Hari et al., 2001; Ninova et al., 2020). Hrb27C is an abundant hnRNP that is involved in various aspects of post-transcriptional mRNA regulation and is implicated in the modulation of Yki activity (Goodrich et al., 2004; Mach et al., 2018). We tested whether these three Bon interactors are involved in trichome formation in adult eyes. Knockdown of *HDAC1* strongly suppressed, whereas its overexpression enhanced, both Bon- and Yki-S168A-induced trichomes (Figures 5B-C, 5G-H, 5L, and S5B-C), and knockdown of *Su(var)2-10* or *Hrb27C* suppressed these trichomes (Figures 5D-E, 5I-J, 5L, and S5B-C). Therefore, HDAC1, Su(var)2-10, and Hrb27C promote Bon- or Yki-induced epidermal fate in the eye.

**Figure 5.**
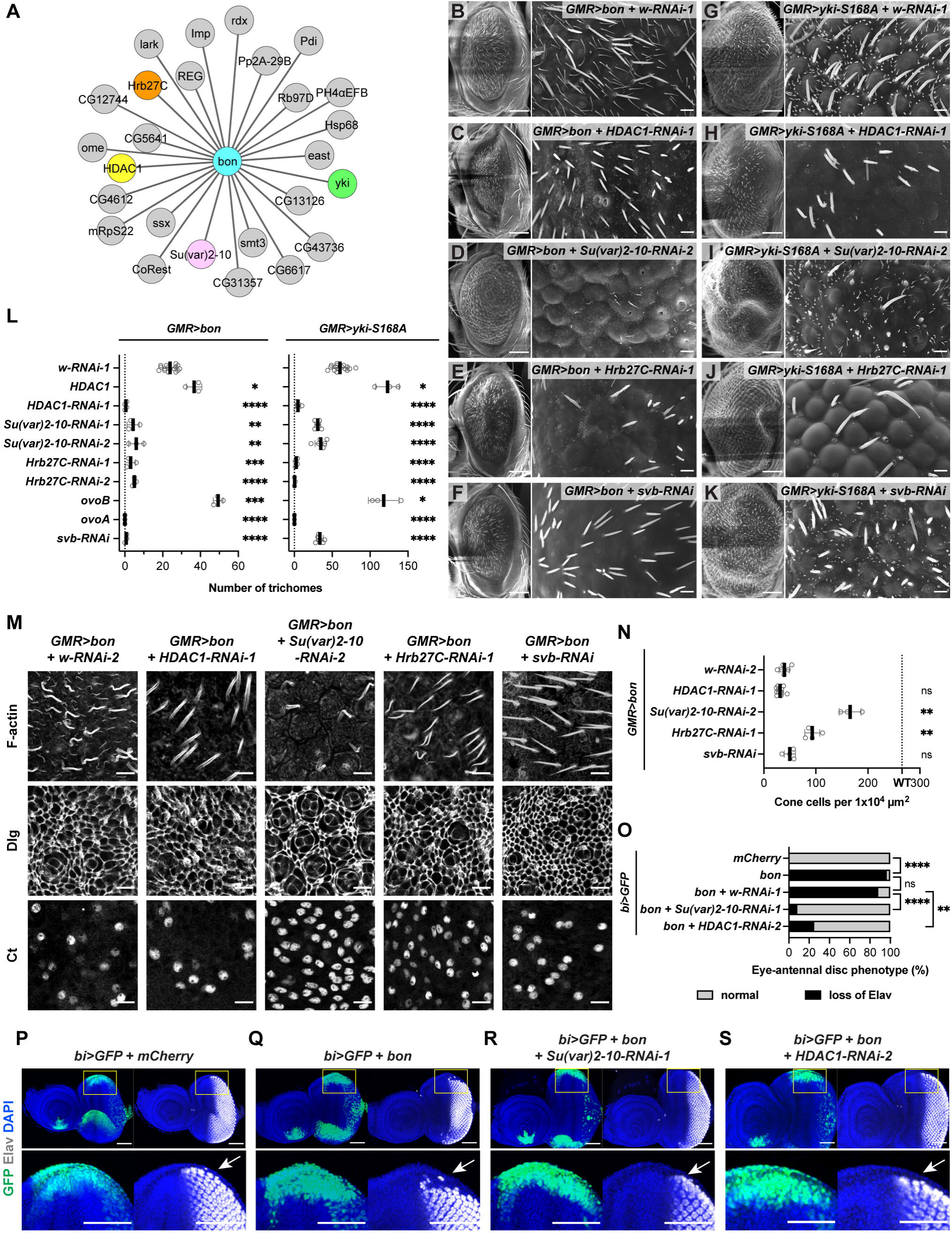
Interactors of Bon and Yki are required for the eye-epidermis fate decision. (A) Bon protein interactome showing significant interactors (SAINT score ≥ 0.8) identified from AP-MS of Bon-SBP in S2 cells. Interactors highlighted with colors were tested in genetic experiments. (B-K) SEM images of adult eyes expressing indicated *UAS* transgenes with the *GMR-GAL4* driver. *w-RNAi-1* was coexpressed for transgene dosage compensation. Crosses with Bon were carried out at 25°C, and crosses with Yki-S168A were set up at 25°C and shifted to 29°C after the emergence of first instar larvae. Scales bars in left panels: 100 μm; enlarged views in right panels: 10 μm. (L) Quantification of the trichome numbers for the indicated genotypes in B-K and S5B-C. (M) Pupal eyes at 44 hrs APF expressing the indicated transgenes with *GMR-GAL4* were stained with phalloidin for F-actin and anti-Dlg antibody for cell boundaries, or anti-Ct antibody for cone cells. Scale bars: 10 μm. (N) Quantification of cone cell numbers per 1×10^4^ μm^2^ area in M. Dashed line indicates the mean cone cell number in WT (*GMR>mCherry*) shown in Figures 3A-B. The numbers are provided in Table S5. (O) Quantification of the phenotypes in eye-antennal discs shown in P-S. P values were determined using Fisher’s exact test between normal and loss-of-Elav populations within indicated comparisons. The numbers are provided in Table S5. (P-S) L3 eye-antennal discs expressing indicated transgenes with the *bi-GAL4* driver were immunostained with anti-Elav antibody. Enlarged views of the dorsal margins (boxed) of the eye discs are shown in the bottom panels. Arrows: Elav expression at the dorsal margins. Scale bars: 50 μm.

The transcription factor Shavenbaby/Ovo (Svb/Ovo) plays a key role in the formation and patterning of epidermal trichomes (Chanut-Delalande et al., 2006; Delon *et al*., 2003). Also, Svb interacts with Yki and regulates apoptosis through *diap1* (Bohere et al., 2018). We tested if Svb/Ovo was involved in the regulation of Bon- and Yki-induced trichomes in adult eyes. Overexpression of the constitutive activator OvoB enhanced Bon and Yki-S168A induced trichomes, while overexpression of the constitutive repressor OvoA, or knockdown of somatic *svb*, strongly suppressed Bon- and Yki-S168A-induced trichomes (Figures 5F, 5K-L, and S5B-C) (Andrews et al., 2000). These results suggest that Svb/Ovo is required for ectopic trichome generation in the eye induced by Bon or Yki.

We then asked if these interactors are involved in Bon and Yki regulated suppression of eye fate. In pupal eyes, knockdown of *HDAC1* or *svb* with *GMR-GAL4* suppressed Bon-induced trichomes but did not rescue the loss of cone cells (Figures 5M-N). In contrast, knockdown of *Su(var)2-10* or *Hrb27C* not only suppressed the trichomes, but also largely rescued the number of cone cells (Figures 5M-N). Interestingly, *Su(var)2-10* RNAi even partially restored the loss of primary pigment cells and mispatterned ommatidial array (Figure S5D). These results suggest that Su(var)2-10 and Hrb27C contribute to both eye fate suppression and epidermal fate promotion in the pupal eye, while HDAC1 and Svb/Ovo are only involved in the latter function at this stage. In L3 eye-antennal discs, Bon overexpression with the *bi-GAL4* driver, which drives expression at the dorsal and ventral margins of the eye disc from L2 stage (Tare et al., 2013), inhibited Elav expression (Figures 5O-Q), similar to Yki overexpression in a previous report (Wittkorn *et al*., 2015). Knockdown of *Su(var)2-10* or *HDAC1* suppressed the loss of Elav due to Bon overexpression (Figures 5R-S and 5O). Furthermore, knockdown of *HDAC1* under *ey-GAL4* rescued the loss of eye field due to Yki-S168A overexpression (Figures S5E-G). Collectively, these results suggest that Bon, Yki, Hrb27C, Su(var)2-10, and HDAC1 work together to suppress eye fate, and promote the epidermal fate with the involvement of Svb/Ovo.

### Bon and Yki control cell fate decisions in the eye through transcriptional regulation of joint target genes

Since Bon, Yki, and interactors described above are all transcriptional or post-transcriptional regulators, we hypothesized that they mediate cell fate choices in the eye-antennal disc via joint regulation of a unique set of transcriptional targets. To identify these target genes, we performed RNA sequencing (RNA-seq) using pupal eyes at 40-41 hrs APF (when the trichomes initiate) that overexpressed Bon, with or without a simultaneous RNAi knockdown of *yki*. Through differential gene expression analysis, we identified 216 genes as Bon-activated/Yki-dependent genes, and 119 genes as Bon-repressed/Yki-dependent genes (Figures 6A, S6A-B, and Table S3). Correlation coefficient analysis revealed the significantly concordant regulation of gene expression by Bon and Yki (Figure 6B). Also, *ex* and *Rhodopsin 5* (*Rh5)*, which are known targets of Yki in the pupal eye (Deng et al., 2020; Jukam *et al*., 2013), were among Yki-dependent Bon-activated genes (Figures 6A and S6A-B). These transcriptomic data suggest that Bon and Yki work together to regulate gene expression in pupal eyes, consistent with our genetic results.

**Figure 6.**
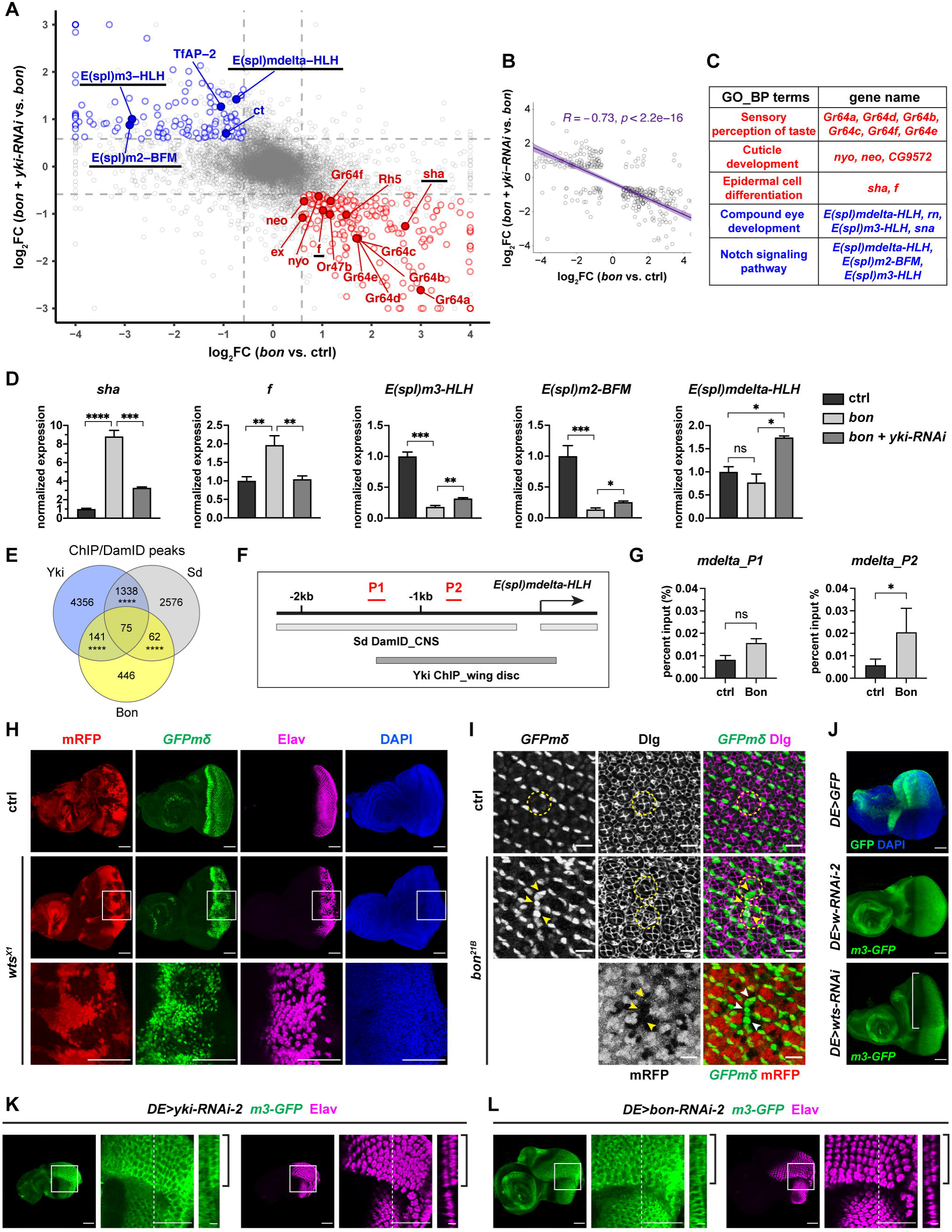
Bon and Yki jointly regulate a set of non-canonical transcriptional target genes. (A) Differential gene expression analysis of RNA-seq data from 40-41 hrs pupal eyes. Output from DEseq2 (Love et al., 2014) was plotted using log_2_ fold changes (FC) for *GMR>bon-mCherry + yki-RNAi-1* vs. *GMR>bon-mCherry* (y axis) and that for *GMR>bon-mCherry* vs. *GMR>mCherry* (WT) (x axis). Dashed lines: FC = 1.5 in both directions. Red circles: Bon-activated/Yki-dependent genes. Blue circles: Bon-repressed/Yki-dependent genes. All colored circles fulfill *p* adj. ≤ 0.05. Solid circles with labels: genes of interest. Labels underlined: genes validated with qRT-PCR. (B) Pearson correlation coefficient analysis of genes jointly regulated by Bon and Yki. Axes are the same as in A. All genes with FC ≥ 1.5 in both directions and *p* adj. ≤ 0.05 were included in the analysis. (C) Terms of interest from GO_BP analysis of Yki-dependent Bon-activated (red) and -repressed (blue) genes. Complete GO_BP analysis is provided in Table S4. (D) qRT-PCR validation for the genes of interest using 40-41 hrs APF pupal eyes with the genotypes *GMR>mCherry* (ctrl), *GMR>bon-mCherry*, and *GMR>bon-mCherry + yki-RNAi-1*. (E) A Venn diagram showing the overlaps of binding sites among Yki, Bon and Sd. All pairwise overlaps have *p* < 0.0001. Previously published datasets used in the analysis were: Yki ChIP-seq from wing discs, Bon ChIP-seq from embryos, and Sd DamID-seq from larval CNS (Negre *et al*., 2011; Oh *et al*., 2013; Vissers *et al*., 2018). (F) Schematic diagram showing a genomic region around the transcription start site (arrow) of *E(spl)mdelta-HLH* (−2.2 kb to +0.5 kb). Yki and Sd binding regions from previous publications are indicated (Oh *et al*., 2013; Vissers *et al*., 2018). P1 and P2: amplicons for ChIP-qPCR. (G) ChIP-qPCR analysis for the direct binding of Bon to the two genomic loci (P1 and P2) shown in F, using L3 eye-antennal discs from *GMR>bon-mCherry*. (H) L3 eye-antennal discs with mosaic clones of the indicated genotypes were immunostained with anti-GFP antibody for *GFPmδ* reporter and anti-Elav antibody for neuronal eye fate. Mutant clones were generated with *eyFLP* and marked by loss of mRFP. Enlarged views of the boxed regions in *wts^X1^* clone are shown in the bottom panels. Scale bars: 50 μm. (I) Pupal eyes at 25 hrs APF of control (*DE>w-RNAi*) and *bon^21B^*mosaic clones were immunostained with anti-GFP antibody for *GFPmδ* reporter and anti-Dlg antibody for cell boundaries. Mutant clones were generated with *eyFLP* and marked by loss of mRFP. Dashed circle: individual ommatidium, arrowheads: ectopic expression of *GFPmδ* in *bon^21B^* clone. Scale bars: 10 μm. (J) L3 eye-antennal discs expressing indicated transgenes with *DE-GAL4*. The expression region of the driver is shown with GFP in the top panel. Signal from *m3-GFP* reporter was amplified by immunostaining with anti-GFP antibody. Bracket: loss of *m3-GFP* in the dorsal compartment of *DE>wts-RNAi*. Scale bars: 50 μm. (K-L) L3 eye-antennal discs expressing indicated transgenes with *DE-GAL4* were immunostained with anti-GFP antibody for *m3-GFP* reporter and anti-Elav antibody for neuronal eye fate. Left and middle panels are focused at the level of endogenous peripodial epithelium (PE). Middle panels: enlarged views of the boxed regions, right panels: orthogonal sections at the dashed lines, brackets: gain of *m3-GFP* and Elav in the PE layer of the dorsal compartment. The orthogonal sections and their scale bars were scaled 2x along the z axis for easier visualization. Scale bars: 5 μm in orthogonal views, 50 μm in others.

We evaluated genes jointly regulated by Bon and Yki by Gene Ontology analysis of Biological Process (GO_BP, Table S4). Bon-activated/Yki-dependent genes were enriched for GO terms “epidermal cell differentiation”, “cuticle development”, and “sensory perception of taste”, while Bon-repressed/Yki-dependent genes were enriched for GO terms “compound eye development” and “Notch signaling pathway”, consistent with the genetic function of Bon and Yki in promoting epidermal fate and antennal fate as well as suppressing the eye fate (Figure 6C). Bon-activated/Yki-dependent epidermal genes, *shavenoid* (*sha*), *forked* (*f*), *nyobe* (*nyo*), and *neyo* (*neo*), are known Svb target genes that are essential for formation and patterning of epidermal trichomes, suggesting that they may mediate Bon- and Yki-induced trichome formation in the eye (Figures 6A and 6C) (Chanut-Delalande *et al*., 2006; Fernandes et al., 2010). Bon-repressed/Yki-dependent Notch pathway genes, *E(spl)mdelta-HLH*, *E(spl)m3-HLH* and *E(spl)m2-BFM*, are members of the Enhancer of split gene complex (*E(spl)-C*), which is a major transcriptional target of Notch (Figures 6A and 6C) (Knust et al., 1992; Lai et al., 2000). Two other Notch target genes, *ct* and *Transcription factor AP-2* (*TfAP-2*), were also identified among Bon-repressed/Yki-dependent genes (Figure 6A) (Kerber et al., 2001; Micchelli et al., 1997). Previous work showed that Notch signaling promotes eye fate and suppresses antennal fate, and that Notch, E(spl)-C, Ct, and TfAP-2 are all critical for cell fate establishment in the eye (Cagan and Ready, 1989b; Kumar and Moses, 2001; Kurata et al., 2000; Lai and Rubin, 2001; Monge et al., 2001; Nagaraj and Banerjee, 2007; Wang and Sun, 2012). These observations raise the intriguing possibility that downregulation of Notch targets by Yki and Bon mediates suppression of eye fate and promotion of antennal fate. Finally, Bon-activated/Yki-dependent genes included sensory perception genes such as antenna-specific odorant receptor *Or47b*, as well as a group of gustatory receptors (*Gr64a-f*) that are expressed in multiple sensory organs including the antenna but excluding the eye, indicating an induction of antennal identity at the transcriptional level (Figures 6A, 6C, and S6A-B) (Barish et al., 2017; Fujii et al., 2015; Menuz et al., 2014; Vosshall, 2001).

We validated the transcriptional regulation of *sha*, *f*, *E(spl)m3-HLH*, *E(spl)m2-BFM*, and *E(spl)mdelta-HLH* by Bon and Yki using target-specific qPCR, which generally confirmed the RNA-seq findings (Figure 6D). To determine whether the transcriptional targets of Yki and Bon are under their direct control, we analyzed published ChIP-seq datasets for Yki and Bon, and the DamID-seq dataset for Sd (Negre et al., 2011; Oh et al., 2013; Vissers et al., 2018). Pairwise overlaps of binding sites were significant, with 75 loci shared among all three proteins, including the *E(spl)-C* region, suggesting a general co-localization of Yki, Bon and Sd on chromosomes and their direct control of *E(spl)-C* (Figure 6E and S6C). Yki and Sd also bound to *ct* and *TfAP-2*, raising the possibility of a direct regulation (Figure S6C). Yki, Sd, and Bon did not associate with the epidermal differentiation genes and antennal sensory receptor genes, except for Yki binding to *neo*, suggesting indirect activation of these genes by Bon and Yki (Figure S6C). To validate the binding of Bon to the *E(spl)mdelta-HLH* genomic locus, we performed ChIP-qPCR using L3 eye-antennal discs expressing Bon-mCherry with *GMR-GAL4*. We chose two DNA segments (P1 and P2) that overlapped the Yki and Sd binding regions (Figure 6F). Bon bound to P2 but did not significantly associate with P1 (Figure 6G). Together, these results suggest that Bon and Yki jointly and directly repress Notch targets and indirectly activate epidermal and antennal markers.

Identification of genes that are directly repressed by Yki and Bon was unexpected and suggests that Bon binding may shift Yki activity towards repression. We further evaluated the repression of *E(spl)-C* by Bon and Yki using a BAC transgene reporter expressing GFP-tagged E(spl)mdelta-HLH (*GFPmδ*) and a transcriptional reporter of *E(spl)m3-HLH* (*m3-GFP*) (Couturier et al., 2019). In the L3 eye disc, both reporters are expressed within and posterior to the MF (Figures 6H and 6J). *wts^X1^* clones showed strongly reduced expression of *GFPmδ* and *m3-GFP*, and moderately reduced Elav (Figures 6H and S6D). In pupal eyes 25 hrs APF, *GFPmδ* is mainly expressed in primary pigment cells (Figure 6I). *bon^21B^* clones resulted in a gain of *GFPmδ*-positive cells, suggesting that Bon suppresses *E(spl)-C* and inhibits the primary pigment cell fate (Figure 6I). *wts* knockdown with the *DE-GAL4* driver, which is expressed in the dorsal eye in both columnar cells of the disc proper (DP) and squamous peripodial epithelium (PE) (Figure S6E) (Morrison and Halder, 2010), reduced *GFPmδ*, *m3-GFP*, and Elav (Figures 6J and S6F). Knockdown of *yki* or *sd* expanded *GFPmδ*, *m3-GFP*, and Elav into the PE (Figures 6K and S6G-I), reminiscent of a PE to DP transformation previously reported for loss of *yki* and *sd* (Zhang *et al*., 2011). *bon* knockdown also led to occasional expansion of *m3-GFP* and Elav into the PE, with a penetrance of 8.3% and 12.5% for two RNAi lines (Figures 6L, S6J, and Table S5). Together, these results show that Bon, Yki and Sd repress, while Wts promotes, the expression of *E(spl)mdelta-HLH* and *E(spl)m3-HLH* in the eye.

We then asked whether the target genes we identified are involved in Bon and Yki control of eye-antenna-epidermal fate determination. Sha overexpression enhanced, whereas *f* knockdown suppressed, trichomes induced by Bon and Yki-S168A in the eye (Figures 7A-C, 7E-G, 7I, S6K-L). Therefore, *sha* and *f* promote Bon/Yki-induced epidermal fate in the eye, consistent with their transcriptional activation by Bon and Yki. In contrast, overexpression of E(spl)mdelta-HLH and E(spl)m3-HLH suppressed the trichomes induced by Bon and Yki-S168A, suggesting that they inhibit the epidermal fate in the eye, in agreement with their transcriptional repression by Bon and Yki (Figures 7D, 7H, 7I, S6M-N). Overexpression of E(spl)m3-HLH significantly rescued the loss of eye field and completely suppressed the formation of double antennae due to Yki-S168A overexpression, suggesting that E(spl)-C promotes eye fate and suppresses antennal fate (Figures 7J-K, S5G, and Table S5). Therefore, Bon and Yki control cell fate decisions in the eye through transcriptional regulation of a unique set of target genes: activation of *sha* and *f*, and repression of *E(spl)-C* genes.

**Figure 7.**
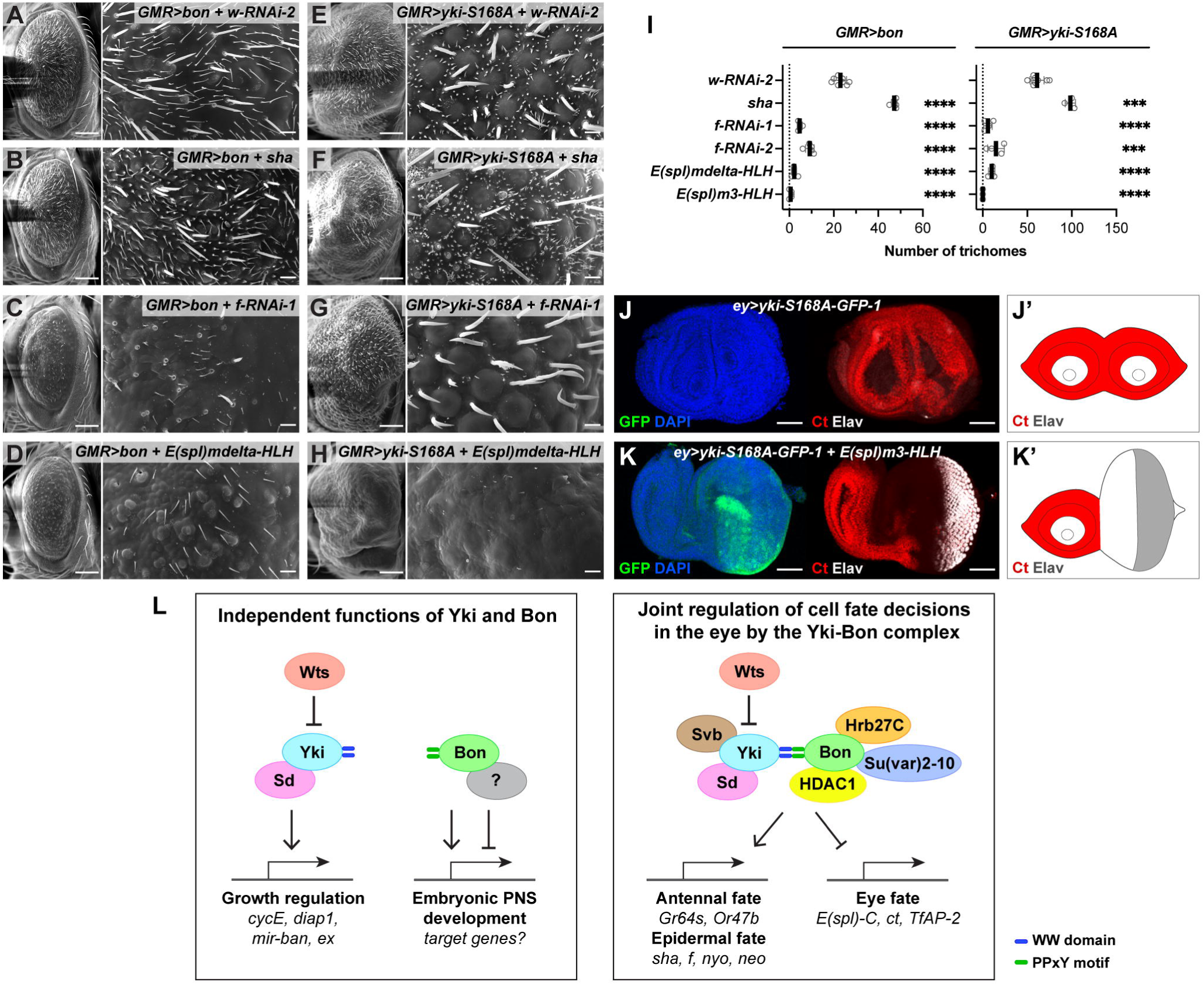
Bon and Yki control the eye-antenna-epidermis fate determination through their joint transcriptional targets. (A-H) SEM images of adult eyes expressing indicated *UAS* transgenes with the *GMR-GAL4* driver. *w-RNAi-2* was coexpressed for transgene dosage compensation. Crosses with Bon were carried out at 25°C, and crosses with Yki-S168A were set up at 25°C and shifted to 29°C after the emergence of first instar larvae. Scales bars in left panels: 100 μm; enlarged views in right panels: 10 μm. (I) Quantification of the trichome numbers for the indicated genotypes in A-H and S6K-N. (J-K’) L3 eye-antennal discs expressing indicated transgenes with *ey-GAL4* were immunostained with anti-Ct and anti-Elav antibodies. (J’-K’) Schematic illustrations of J-K. Scale bars: 50 μm. Quantification of the phenotypes is shown in S5G, and the detailed numbers are given in Table S5. (L) Left: independent functions of Yki and Bon in growth regulation and embryonic PNS development. Right: Yki and Bon jointly regulate cell fate decisions in the eye through recruitment of transcriptional and post-transcriptional regulators and transcriptional control of a set of non-canonical target genes.

## Discussion

We have revealed a previously unappreciated and profound role of the Hippo pathway in controlling cell fate determination in the *Drosophila* eye (Figure 7L). This function depends on the activity of the novel Yki interactor, Bon. We have uncovered a collaborative regulatory program in which Yki and Bon interact, and likely function within larger multiprotein complexes that include other transcriptional regulators. Rather than controlling growth or PNS differentiation, which represent the previously ascribed independent functions of Yki and Bon, the Yki-Bon module regulates the proper segregation of the eye, epidermal, and antennal fates in the developing eye. This function involves the promotion of the epidermal and antennal fates, and the suppression of the eye fate, via transcriptional regulation of a distinct set of target genes (Figure 7L). Our study thus provides a molecular mechanism for the biological function of the Hippo pathway and Bon in cell fate determination during eye development.

### Hippo pathway and Bon control eye-antenna-epidermis cell fate decisions at two levels

Our results suggest that the Hippo pathway and Bon regulate the developmental cell fate decisions in the eye at two levels (Figure S7). First, the Yki-Bon complex promotes antennal and epidermal fates and suppresses the eye fate during the early eye field specification, before the L3 larval stage. This is supported by the phenotypes observed under various genetic manipulations of Bon, Wts, Yki, and Sd during the L1 and L2 larval stages, including the reciprocal transformations of eye and antenna, epidermal outgrowths in the eye, and ectopic eye fate (Figures 3, 4, 6, S4, and S6). The Yki-Bon module is thus an essential component of the extensive network of signaling pathways and transcription factors that control these cell fate decisions in early eye development (Kumar and Moses, 2001). Previous studies showed that *ex*, *mer* and *mats* mutants exhibit eye to epidermal transformation and occasionally eye to antenna transformation (in *ex* mutants), suggesting that the upstream Hippo pathway may also regulate the Yki-Bon controlled fate determination at this stage (Boedigheimer and Laughon, 1993; Lai *et al*., 2005; McCartney *et al*., 2000; Pellock *et al*., 2007).

Second, after the segregation of the eye/antenna/epidermis fields and the start of MF in L3, the Yki-Bon complex promotes the epidermal cell fate while suppressing ommatidia, whereas Wts counteracts this activity. This is evidenced by the formation of epidermal trichomes on interommatidial cells as well as suppression of ommatidial cell types, especially cone cells, upon knockdown of *wts* or overexpression of Bon or Yki with the late eye driver *GMR-GAL4* (Figures 2, 3, and S3). Furthermore, our RNA-seq data also revealed cell fate regulation at the gene expression level: the Yki-Bon complex activates epidermal differentiation genes (*sha*, *f*, *nyo* and *neo*), and represses multiple Notch targets (*E(spl)-C*, *ct* and *TfAP-2*) that are required for eye fate establishment and are expressed in ommatidial cells (e.g., *ct* in cone cells and *E(spl)mdelta-HLH* in primary pigment cells) (Figure 6). Although activation of the Yki-Bon complex at this stage did not exhibit an eye-to-antenna transformation phenotype, certain antennal genes (*Gr64s* and *Or47b*) were upregulated, suggesting transformation at the level of gene expression.

The eye-antenna-epidermis fate determination was previously studied during the early stages of eye-antennal disc development (Kumar, 2018). Our work shows that these fates are not completely defined during the early stages, as the retina could still transform into epidermal tissue and express epidermal and even antennal genes when the Hippo pathway and Bon were modulated after MF formation. Interestingly, conditional knockout of *eya* after the MF results in suppression of ommatidia and formation of trichomes in the eye (Jin et al., 2016). This suggests that the retinal determination genes are also involved in eye-epidermal fate decisions during later stages of eye development, and that trichome induction may be a general biological outcome of interference with the eye vs. epidermis specification after the start of MF. Thus, we conclude that the eye is still developmentally plastic at late stages, with a latent epidermal fate that is normally inhibited. Interestingly, this fate is revealed in the insect order Strepsiptera, whose compound eyes are comprised of optical units that are separated by epidermal tissue bearing trichome-like extensions (Buschbeck, 2005).

### Yki and Bon control cell fate decisions in the eye by recruiting cofactors and regulating a distinct set of target genes

We have uncovered an unexpected layer of control over eye specification exerted by Yki and Bon at the level of Notch target genes. The Hippo pathway has been reported to control cell fate determination in other biological contexts through regulation of Notch ligands (Reddy et al., 2010; Totaro et al., 2017). Although we have identified several Notch targets that are repressed by Bon and Yki, Notch itself and the key Notch pathway components, Serrate, Delta, and Su(H), were not transcriptionally regulated or found in Yki or Bon protein interactomes (Tables S1-S3). Therefore, we propose that during cell fate determination in the eye, Bon and Yki repress Notch targets such as *E(spl)-C* genes independently from upstream Notch signaling. We note that not all *E(spl)-C* genes are under Bon and Yki control, implying context-dependent regulation and functional divergence of *E(spl)-C* genes, as suggested by previous studies (Table S3) (Couturier *et al*., 2019; Ligoxygakis et al., 1999). Both Hippo and Notch contribute to cell proliferation and growth of the eye (Dominguez and Casares, 2005; Peng et al., 2009). Our data suggest that Bon is not required for the growth regulation function of the Hippo pathway (Figure S1). Instead, the interaction between Bon and Yki is specifically required for controlling eye specification through regulation of Notch targets. We speculate that Bon may function as a switch that redirects some of Hippo pathway activities from growth regulation to cell fate determination.

So far, *Drosophila* Yki has only been implicated in transcriptional activation (Zheng and Pan, 2019). However, studies in mammalian systems have shown that the YAP/TAZ-TEAD complex (Yki-Sd orthologs) can also function as a transcriptional repressor on non-canonical target genes (Beyer et al., 2013; He et al., 2020; Kim et al., 2015). The direct repression of Notch targets reported here suggests that *Drosophila* Yki can also function in transcriptional repression, likely via recruitment of corepressors mediated by Bon. HDAC1 and its associated corepressor complexes repress gene transcription (including Notch targets) via epigenetic modifications (Borggrefe and Oswald, 2009). We identified HDAC1 and its corepressor, CoRest, in the Bon interactome (Figure 5A), raising the possibility that Bon and Yki repress Notch target genes in part via recruitment of this repressor complex. The involvement of epigenetic regulators is further exemplified by Su(var)2-10, which has a role in chromatin SUMOylation and piRNA target gene silencing (Ninova *et al*., 2020). Interestingly, Su(var)2-10 can suppress eye fate and even induce antennal fate in a sensitized background (Betz *et al*., 2001). Due to the strong genetic interaction between Su(var)2-10 and the Bon-Yki complex (Figures 5 and S5), it is possible that chromatin SUMOylation is involved in gene repression by Bon and Yki. Future studies of chromatin status and epigenetic markers may reveal the mechanistic details of gene repression by Bon and Yki.

The Hippo pathway and Bon orthologs are conserved and broadly expressed in higher eukaryotes (Cammas *et al*., 2012; Zheng and Pan, 2019), raising the possibility that they may also function together in other species and tissue contexts. The vertebrate Hippo pathway controls retinogenesis and the cell fate decision between the neuronal retina and non-neuronal retinal pigment epithelium, which shares similarities with *Drosophila* interommatidial cells (Charlton-Perkins and Cook, 2010; Lee et al., 2018). It is possible that the TIF1 family proteins are also involved in this process. The Hippo pathway and TIF1γ have been independently implicated in controlling hematopoiesis (Lundin et al., 2020; Milton et al., 2014; Rossmann et al., 2021). The YAP/TAZ-TEAD complex or TIF1β can recruit the transcriptional corepressor NuRD complex for gene repression in various biological contexts (Beyer *et al*., 2013; He *et al*., 2020; Kim *et al*., 2015; Schultz et al., 2001). These studies and our data suggest that the biological functions controlled by the Hippo pathway and Bon and the underlying molecular mechanism we report here may be evolutionarily conserved.

## Acknowledgments

We thank Spyros Artavanis-Tsakonas, Hugo Bellen, Kwang-Wook Choi, Claude Desplan, Maxim Frolov, Jin Jiang, Eric Lai, Duojia Pan, François Schweisguth, Alexei Tulin, BDSC, VDRC, DGRC and DSHB for fly stocks and reagents. We thank Georg Halder, Kieran Harvey, Tony Ip, Kenneth Irvine, Ruth Johnson, Donald Ready, Nicolas Tapon, Barry Thompson, Can Zhang and Lei Zhang for helpful discussions. We thank Alexander Letizia and Katerina Heath for helping with plasmid construction. We also thank Jens Rister and Claire Jackan for helpful comments on the manuscript. Part of the SEM work was supported by Award #S10RR021043 from the National Center for Research Resources to the UMass Chan EM Core Facility. This work was funded by NIH grants GM105813 and GM123136 to K.H.M. and A.V.

## Author contributions

Conceptualization, H.Z., K.H.M., and A.V.; Methodology, H.Z., K.H.M., and A.V.; Formal analysis, H.Z.; Investigation, H.Z.; Resources, K.H.M. and A.V.; Data Curation, H.Z.; Writing – Original Draft, H.Z.; Writing – Review & Editing, H.Z., K.H.M., and A.V.; Visualization, H.Z.; Supervision, A.V.; Funding Acquisition, K.H.M. and A.V.

## Methods

### Plasmid construction

pMK33-yki-SBP was generated by cloning Yki-RG isoform from pMT-yki-V5 (Zhang *et al*., 2015) into pMK33-SBP-C vector (Yang and Veraksa, 2017). pUASTattB-yki-EGFP was generated by cloning Yki-RG and EGFP tag into pUASTattB vector (Bischof et al., 2007). pMT-yki-HA was generated by cloning Yki-RG and HA tag into pMT-V5-His vector (Invitrogen, Cat# V412020). The Y281A, Y350A mutations of WW domains in Yki were generated by Q5 Site-Directed Mutagenesis Kit (NEB, Cat# E0554S) using pMT-yki-HA as a template. The numbering of the key tyrosine residues used in this work corresponds with the first reported yki sequence (yki-RH) which has a stretch of additional 23 aa at the N-terminus (Huang *et al*., 2005). All tags were added to the C-terminus of the genes.

pMT-bon-V5, pMK33-bon-SBP, and pUASTattB-bon-mCherry were generated by cloning Bon-RB isoform from pFlc-1-bon (DGRC, RE48191). The Y507A, Y585A mutations in PPxY motifs of Bon were generated by the Q5 Site-Directed Mutagenesis Kit. pUASTattB-bon-mCherry and pUASTattB-bon-Y507A Y585A-mCherry were constructed using NEBuilder HiFi DNA Assembly Cloning Kit (NEB, Cat# E5520S). All tags were added to the C-terminus of the genes.

pMT-Myc-wts was generated by cloning the coding sequence of wts from pFlag-wts plasmid kindly offered by Maxim Frolov Lab. Myc tag was added to the N-terminus.

### Stable cell lines

S2 cells were transfected with pMK33-yki-SBP and pMK33-bon-SBP constructs using the Effectene transfection reagent (Qiagen, Cat# 301427). Stable cell lines were selected with medium containing 300 μg/ml of Hygromycin B (MilliporeSigma, Cat# H3274). Yki-SBP and Bon-SBP stable cells were induced overnight with 0.07 mM CuSO_4_, and protein expression was verified by western blot using anti-SBP antibody (1:1000, Santa Cruz, Cat# sc-101595).

### Transgenic fly lines

pUASTattB-yki-EGFP plasmid was injected into the attP2 site in *D. melanogaster* embryos (Genetic Services, Inc). pUASTattB-bon-mCherry and pUASTattB-bon-Y507A Y585A-mCherry plasmids were injected into the attP40 site in *D. melanogaster* embryos (Rainbow Transgenic Flies, Inc). *yw*, Chr. 2 and Chr. 3 balancer stocks were used in standard crossing schemes to establish the homozygous transgenic lines.

### Genetic crosses

To score the trichome phenotype, *UAS-bon-mCherry*, *UAS-yki-EGFP*, *UAS-yki-S168A-1* (*UAS-yki.S168A.GFP.HA*, BDSC: 28816) and *UAS-wts*-RNAi (VDRC: 111002) were combined with the eye driver *GMR-GAL4* to generate *UAS-bon-mCherry/CyO wg-lacZ; GMR-GAL4/TM6B*, *GMR-GAL4 UAS-yki-EGFP/TM6B*, *GMR-GAL4 UAS-yki-S168A-GFP-1/TM6B* and *UAS-wts*-RNAi*/CyO wg-lacZ; GMR-GAL4/TM6B* fly lines, respectively. *GFP*, *w*-RNAi-1, or *w*-RNAi-2 were coexpressed for transgene dosage compensation.

To achieve higher GAL4 activity, all crosses with *GMR-GAL4 UAS-yki-EGFP*, *GMR-GAL4 UAS-yki-S168A-GFP-1*, and *UAS-wts*-RNAi *GMR-GAL4* were set up at 25°C and shifted to 29°C after the emergence of first instar larvae. All the crosses with *UAS-bon-mCherry/CyO wg-lacZ; GMR-GAL4/TM6B* were carried out at 25°C.

All the crosses using *ey-GAL4* and *ey-GAL4-2* were carried out at 29°C except for the cross of *ey-GAL4* and *UAS-wts*-RNAi which was carried out at 25°C. The cross of *da-GAL4* and *UAS-yki-EGFP* for embryo collection and affinity purification was carried out at RT. All other crosses were carried out at 25°C.

Mosaic clones with hsFLP/FRT were generated by heat-shocking first instar larvae with indicated time after egg deposition (AED) at 37°C for 1 hr. Genotypes used for clones are as follow:

3E: hsFLP;; FRT82B/FRT82B Ubi-GFP

3F-G: hsFLP;; FRT82B wts^X1^/FRT82B Ubi-GFP

3H: hsFLP;; FRT82B wts^X1^/FRT82B Sb63b

3I ctrl: eyFLP;; FRT82B/FRT82B Ubi-GFP

3I bon^21B^: eyFLP;; FRT82B bon^21B^/FRT82B Ubi-GFP

6H ctrl: eyFLP;; GFPmδ/+; FRT82B/FRT82B Ubi-mRFP

6H wts^X1^: eyFLP;; GFPmδ/+; FRT82B wts^X1^/FRT82B Ubi-mRFP

6I bon^21B^: eyFLP;; GFPmδ/+; FRT82B bon^21B^/FRT82B Ubi-mRFP

S6D ctrl: *eyFLP;; m3-GFP/+; FRT82B/FRT82B Ubi-mRFP*

S6D *wts^X1^*: *eyFLP;; m3-GFP/+; FRT82B wts^X1^/FRT82B Ubi-mRFP*

### Affinity Purification

#### Affinity purification from S2 cells

Blank S2 cells (control) and stable cell lines with 50 ml dense culture were induced overnight (∼16 hrs) with 0.07 mM CuSO_4_ at 25°C. Cells were collected and lysed on ice for 20 min using 1 ml of ice-cold Default Lysis Buffer (DLB) (50 mM Tris pH 7.5, 125 mM NaCl, 5% (v/v) glycerol, 0.2% (v/v) IGEPAL, 1.5 mM MgCl_2_, 25 mM NaF, 1 mM Na_3_VO_4_, 1 mM DTT, and 2x cOmplete Protease Inhibitor (MilliporeSigma, Cat# 11836145001, 1 tablet per 25 ml lysis buffer)). The lysate was centrifugated at 16,000 rcf for 20 min at 4°C, and the supernatant was incubated with 50 μl of packed Streptavidin beads (Thermo Fisher Scientific, Cat# 53117) for 2 hrs at 4°C with rotation. The beads were then washed 5 times with 1 ml DLB. Proteins on the beads were eluted with 300 μl DLB containing 2 mM biotin and precipitated by adding 1/10 volume of 8.74 M Trichloroacetic acid solution (TCA, Fisher Scientific, Cat# BP555-500). Precipitated proteins were washed once with 500 μl of 0.874 M TCA and four times with ice-cold acetone. The protein pellet was dried overnight at RT and incubated in 40 μl of 2x SDS sample buffer at 95°C for 5 min. Samples were assessed by silver staining using NuPAGE 4-12% Bis-Tris Protein Gel (Thermo Fisher Scientific, Cat# NP0335) and SilverQuest Silver Staining Kit (Thermo Fisher Scientific, Cat# LC6070). The protein samples for mass spectrometry were separated on a short SDS-PAGE gel (8% Tris-Glycine) followed by Coomassie blue staining. Two gel pieces (>75 kDa and < 75 kDa) for each sample were submitted to Taplin Mass Spectrometry Facility at Harvard Medical School for mass spectrometry analysis, as described (Yang and Veraksa, 2017).

#### Affinity purification from embryos

5 L fly cages with apple juice plates (Yang et al., 2017) were set up for *yw* (control) and the cross *da-GAL4* x *UAS-yki-EGFP*. Flies were allowed to lay eggs for 15 hrs at RT, then the apple juice plates containing the embryos were incubated for 3 hrs at 25°C. Approximately 1 g embryos were dechorionated with 50% (v/v) Clorox bleach and washed with water. Collected embryos were then mixed with 6 ml of ice-cold DLB containing 0.5% (v/v) IGEPAL and 10 μM MG132 in a glass homogenizer and lysed on ice using 20-30 strokes with a tight pestle. The lysate was kept on ice for 20 min and centrifuged at 16,000 rcf for 20 min at 4°C. The supernatant was filtered with pre-chilled 0.45 μm filter and incubated with 50 μl of packed Pierce Control Agarose Resin (Thermo Fisher Scientific, Cat# 26150) for 30 min at 4°C with rotation to minimize unspecific binding. The lysate was then incubated with 20 μl of packed GFP-Trap Agarose (Bulldog Bio, Cat# GTA020) for 2 hrs at 4°C with rotation. After binding, the GFP beads were washed 5 times with 1 ml DLB containing 0.5% (v/v) IGEPAL and 10 μM MG132, followed by addition of 40 μl of 4x SDS sample buffer and heating at 95°C for 6 min. The samples were then assessed by silver staining and the protein samples for mass spectrometry were prepared and submitted as above.

### Mass Spectrometry and data analysis

In-gel trypsin digestion and liquid chromatography/tandem mass spectrometry (nanoLC-MS/MS) were performed by Taplin Mass Spectrometry Facility at Harvard Medical School. Mass spectrometry results from two gel pieces for each sample were combined, and the results from the experimental and control samples were analyzed with Significance Analysis of INTeractome (SAINT) (v2.5.0) using the total peptides identified for each protein (Choi *et al*., 2011). Any protein with the SAINT score (average probability) ≥ 0.8 was considered as a high-confidence interactor and was included in the interactome. SAINT analysis included the following numbers of samples: Yki-SBP, four experiments and four controls; Yki-EGFP, three experiments and five controls; Bon-SBP, two experiments and three controls.

#### Yki protein interactome

Yki protein interactors with SAINT score ≥ 0.8 either in S2 cells or in embryos were included in the Yki protein interactome. The Yki protein interactome was further populated with the interactions incorporated from the STRING database (v11.0) and FlyBase (vFB2020_04) (Szklarczyk *et al*., 2019; Thurmond *et al*., 2019). All proteins in the Yki interactome from the current study were imported into the STRING database and analyzed with default settings: full network, medium confidence 0.4 and all active interaction sources (Textmining, Experiments, Databases, Co-expression, Neighborhood, Gene Fusion and Co-occurrence). The summary of physical interactions of Yki in FlyBase was selected to show neighbor-neighbor interactions, and the summary of genetic interactions was selected to show both suppression and enhancement. The interaction data from FlyBase were exported through esyN (Bean et al., 2014), while only the interactions between the proteins that were identified as components of the Yki protein interactome from the current study were incorporated into the Yki protein interactome. The Yki protein interactome was visualized with Cytoscape (Shannon et al., 2003). Nodes represent Yki and Yki interactors identified in the current study. Edges represent the interactions incorporated from STRING and FlyBase. Clustering was done manually based on FlyBase annotations and publications (Thurmond *et al*., 2019). Gene symbols were updated to FlyBase version FB2020_04, released Aug 18, 2020.

#### Bon protein interactome

Bon protein interactors with SAINT score ≥ 0.8 from S2 cells were included in the Bon protein interactome which was visualized with Cytoscape. The nodes and edges represent the interactors and interactions identified in the current study, respectively. Gene symbols were updated to FlyBase version FB2020_04, released Aug 18, 2020.

### Co-Immunoprecipitation

S2 cells were transfected with indicated plasmids or blank pMT-V5-His vector (Invitrogen, Cat# V412020) using the Effectene transfection reagent (Qiagen, Cat# 301427). 24 hrs after transfection, cells were induced with 0.35 mM CuSO_4_ overnight at 25°C, collected and lysed on ice for 20 min with 600 μl of ice-cold DLB. The lysate was centrifuged at 16,000 rcf for 20 min at 4°C. 40 µl supernatant was mixed with 20 μl of 4x SDS sample buffer and heated at 95°C for 6 min to generate lysate samples. The rest of the lysate was incubated with 20 μl of packed anti-V5 beads (MilliporeSigma, Cat# A7345) or anti-HA beads (MilliporeSigma, Cat# E6779) for 2 hrs at 4°C. The beads were then washed 3 times (or 4 times for co-IP with Myc-wts) with 1 ml DLB, mixed with 40 μl of 4x SDS sample buffer, and heated at 95°C for 6 min to generate IP samples. The lysate samples and IP samples were subjected to SDS-PAGE followed by western blot using Odyssey Blocking Buffer (PBS) (LI-COR Biosciences, Cat# 927-40003) as blocking buffer, mouse anti-V5 antibody (1:1000, MilliporeSigma, Cat# V8012), Rabbit anti-HA antibody (1:1000, MilliporeSigma, Cat# H6908) and Mouse anti-Myc antibody (1:1000, Cell Signaling, Cat# 2276S) as primary antibody, and Goat anti-Mouse IgG (1:20,000, LI-COR Biosciences, Cat# 926-68070) and Donkey anti-Rabbit IgG (1:20,000, LI-COR Biosciences, Cat# 926-32213) as secondary antibody. Blotting membranes were scanned with LI-COR Odyssey CLx Imaging Systems.

### EdU incorporation assay

EdU incorporation assay was performed using Click-iT® EdU Imaging Kit (Thermo Fisher Scientific, C10086) with adapted procedures from (Gouge and Christensen, 2010). Eye discs were dissected in unsupplemented Schneider’s *Drosophila* Medium and incubated in 100 μM EdU for 20 min in the dark at RT. Discs were then washed three times with 1x PBS, fixed in 3.7% (v/v) Formaldehyde (MilliporeSigma, Cat# F8775) in 1x PBS for 15 min in the dark at RT, followed by three washes with 1x PBS and permeabilization in PB-Triton (1x PBS containing 0.3% (v/v) Triton X-100) for 20 min at RT. Discs were then washed twice with 3% BSA in 1x PBS and incubated in Click-iT reaction cocktail containing Alexa Fluor 488 (for *bon-mChery*) or 594 (for *yki-S168A-GFP*) for 30 min in the dark at RT. After washing once with 3% BSA in 1x PBS and once with 1x PBS, discs were mounted in ProLong Gold Antifade Mountant with DAPI (Thermo Fisher Scientific, Cat# P36931). All Incubation and washing were carried out on a platform shaker with mild shaking.

### Immunostaining, fluorescence and bright-field microscopy

#### Antibodies and phalloidin

All antibodies and phalloidin reagents used in this study are listed in the Key Resources Table. All primary antibodies from DSHB were used at 1:50 (v/v) dilution. All primary and secondary antibodies and phalloidin from Thermo Fisher Scientific were used at 1:500 (v/v) dilution. Mouse anti-β-Gal antibody (Promega, Cat# Z3783) was used at 1:100 (v/v) dilution. Guinea Pig anti-Bon antibody was used at 1: 5000 (v/v) dilution (Beckstead *et al*., 2001). Guinea Pig anti-Hth antibody was used at 1:1000 (v/v) dilution (Ozel et al., 2021). Rabbit anti-BarH1 was used at 1:500 (v/v) dilution (Kang et al., 2014). Phalloidin 405 (Biotium, Cat# 00034-T) was used at 1:100 (v/v) dilution.

#### Embryo staining

Stage 16 embryos were collected and dechorionated with 50% (v/v) Clorox bleach, rinsed with water, and fixed in 20 ml of fixative containing 4% (v/v) paraformaldehyde (Electron Microscopy Sciences, Cat# 15710), 25% (v/v) Heptane and 1x PBS for 20 min at RT. 8 ml of methanol was added to remove the vitelline membrane. Fixed and devitellinized embryos were washed three times with 1.4 ml methanol, four times with 1.4 ml ethanol and twice with PBT (1x PBS containing 0.1% (v/v) Tween 20). Embryos were subsequently incubated in 1 ml of blocking solution (1:1 (v/v) of Western Blocking Reagent (Roche, Cat#11921681001) and PBT) for 2 hrs at RT and incubated overnight in 500 μl of blocking solution containing 22C10 primary antibody (DSHB, Cat# 22c10) at 4°C. Embryos were then washed five times with 0.1% BSA (m/v in PBT) and incubated in 1 ml of blocking solution for 1 hr at RT. The embryos were then incubated in secondary antibody diluted in blocking solution for 2 hrs in the dark at RT, washed sequentially in 0.1% BSA, PBT and 1x PBS, and mounted with ProLong Gold Antifade Mountant with DAPI. All incubation and washing were carried out on a nutator.

#### L3 imaginal disc staining

3^rd^ instar larval wing discs and eye-antennal discs were dissected in ice-cold 1x PBS and were fixed in 3.7% (v/v) formaldehyde for 15-20 min at RT. The discs were washed three times with 1x PBS, permeabilized with PB-Triton for 20 min at RT and were incubated in primary antibodies diluted in blocking solution (1:1 (v/v) of Western Blocking Reagent and PBT) overnight at RT. The discs were then washed four times with PBT, incubated in secondary antibodies diluted in blocking solution for 2-3 hrs in the dark at RT, washed four times with PBT, and mounted with ProLong Gold Antifade Mountant with DAPI. All incubation and washing were carried out on a platform shaker with mild shaking.

#### Pupal eye staining

Pupal eye staging, dissection and staining procedures were adapted from (Hsiao et al., 2012; Walther and Pichaud, 2006). Newly formed white pupae were circled on the fly vial. 32-46 hrs after puparium formation (APF), pupal eyes were dissected with forceps in ice-cold 1x PBS, transferred to 3.7% (v/v) formaldehyde (MilliporeSigma, Cat# F8775) and fixed for 15-20 min at RT. Fixative was replaced with 1x PBS and washed three times followed by permeabilizing with PB-Triton for 30 min. The pupal eyes were then incubated in primary antibodies diluted in PB-Triton overnight at RT, washed four times with PB-Triton, and incubated in secondary antibody or Phalloidin diluted in PB-Triton for 3 hrs in the dark at RT. The pupal eyes were then washed four times with PB-Triton and mounted with ProLong Gold Antifade Mountant (Thermo Fisher Scientific, Cat# P10144). All incubation and washing were carried out on a platform shaker with mild shaking.

#### Microscopy

Fluorescent images were acquired using Zeiss LSM 880 Confocal Microscope. For embryos, 20x objective and 1 Airy Unit (AU) pinhole were used. For L3 wing discs, 20x objective was used to take a z stack, and maximum intensity projection of the entire disc proper was performed to generate final images. For L3 eye-antennal discs, 20x or 63x objective was used to take z stacks with optimal step size, and maximum intensity projection was performed for the entire disc proper, except for the anti-Ct staining which was focused on the disc proper while minimizing the inclusion of the cone cell focal plane, or for the orthogonal sections and their corresponding 2D views which are indicated in the figures and legends. For pupal eyes, 63x objective and z stack were used with focal plane set to the apical surface, except for the final images of *GFPmδ* and anti-Elav staining which were generated by maximum intensity projection of the entire pupal eye unless indicated otherwise. In mispatterned pupal eyes overexpressing Bon, Ct-positive cone cells and BarH1-positive primary pigment cells were distinguished from bristle groups which have both Ct and BarH1 expression in the basal nuclei (Beam and Moberg, 2010; Cadigan and Nusse, 1996; Charlton-Perkins and Cook, 2010). Bright-field images of adult eyes were taken under Zeiss Stemi 2000-C Microscope with an attached-on camera at 50x magnification. Due to the difference in size, only images from female adult flies were analyzed. Images were processed with Fiji (Schindelin et al., 2012).

### Scanning electron microscopy (SEM)

Preparation of adult flies for SEM was adapted from (Pepple et al., 2007; Wolff, 2011). 1-2 days old adult flies were sequentially dehydrated in 1 ml of 25%, 50%, 75%, 100%, 100% and 100% ethanol (v/v in water) for 8-16 hrs per step at RT (overnight for the first step). The flies were then chemically dried in 500 μl of 25%, 50%, 75%, 100%, 100% and 100% Hexamethyldisilazane (v/v in ethanol) (HMDS, Electron Microscopy Sciences, Cat# 16700) for 30 min per step at RT. Most of the HMDS was then removed and the remaining HMDS was allowed to be dried under vacuum in a desiccator containing Indicating Drierite (W.A. Hammond, Cat# 23005) overnight at RT. The samples were then mounted on Aluminum Mount using Carbon Adhesive Tape, and stored in Mount Holder (Electron Microscopy Sciences, Cat# 75220, 77816 and 76510). The samples were subsequently coated with gold at 50 mAmps for 150 secs with Denton Vacuum Desk V sputter coater or coated with platinum for 8 nm with Denton Vacuum 502-B for samples in Figures S2O-T. Most images were acquired with JEOL JSM-6010LA IntouchScope Scanning Electron Microscope using 20 kV voltage, 10 mm working distance, 40-50 spot size, and 150x, 300x and 1000x magnifications for the whole eye, zoomed-in view for ectopic antenna, and zoomed-in view for trichomes, respectively. Images in Figures S2O-T were acquired with FEI Quanta 200 FEG MK II Scanning Electron Microscope using 10 kV voltage, 10 mm working distance, 2.5 spot size, and 150x and 1000x magnifications for the whole eye and zoomed-in view, respectively. Due to the difference in size, only images from female adult flies were analyzed. Images were processed with Fiji. The numbers of trichomes were counted with Fiji in an area of 1306 μm^2^ (41.73 μm x 31.30 μm) per image for all genotypes.

### Total RNA preparation

Total RNA preparation method was adapted from (Zhang *et al*., 2015). Pupal eyes were dissected from 40-41 hrs APF pupae (newly formed white pupae were staged for 40 hrs at 25°C and dissected within 1 hr at RT) in ice cold RNase-free 1x PBS diluted from RNase-free 10x PBS (Fisher Scientific, Cat# BP3994) with RNase-free water (Thermo Fisher Scientific, Cat# 10977015). Dissected pupal eyes were immediately transferred to 100 μl of TRIzol (Thermo Fisher Scientific, Cat# 15596026) and kept at RT for 5 min to lyse. Samples in TRIzol were stored at −80°C until 60 pupal eyes for each genotype were collected. All TRIzol samples for each genotype were then thawed and combined, and the volume was brought up to 500 μl per sample. Total RNA was extracted twice by adding 100 μl of chloroform, vigorously shaking for 15 secs, incubating at RT for 3 min, centrifuging at 11,000 rcf at 4°C for 15 min, and collecting the aqueous phase. The total RNA was then precipitated by adding 1 volume of 70% ethanol (v/v in RNase-free water), and purified with RNeasy Mini Kit (QIAGEN, Cat# 74104, 79254).

### RNA-sequencing, data analysis, and qPCR validation

The RNA-seq libraries were prepared from 100 μg of total RNA per genotype with NEBNext Ultra II RNA Library Prep Kit mRNA Isolation Module (NEB, Cat# E7775S, E7490S, E7500S). Quality control and sequencing of the RNA-seq libraries were performed by GENEWIZ, Inc. using Agilent TapeStation and Illumina HiSeq 4000 with a 2×150 paired-end configuration.

The adapter sequences were trimmed from raw reads, and reads shorter than 18 nt were removed by Cutadapt (v2.9) (Martin, 2011). The reads mapped to tRNAs, rRNA, snRNA and snoRNA were removed by mapping through Bowtie 2 (v2.3.5.1) with very sensitive setting and maximum fragment size of 1000 in addition to default setting (Langmead and Salzberg, 2012). The tRNA, rRNA, snRNA and snoRNA reference sequences were obtained from FlyBase (dmel-all-tRNA-r6.33.fasta and dmel-all-miscRNA-r6.33.fasta) (Thurmond *et al*., 2019). The remaining reads were mapped to *D. melanogaster* genome Ensembl_BDGP6 (Flicek et al., 2014) by STAR (v2.7.0e) (Dobin et al., 2013) with default parameters. Gene counting was achieved by featureCounts (Subread v1.6.2) with overlapping or polycistronic genes counted as a fraction (Liao et al., 2014). Differential gene expression analysis was performed with DEseq2 (v1.30.1) through DEBrowser (v1.16.3) with default settings (Kucukural et al., 2019; Love *et al*., 2014), and genes with maximum count fewer than 10 were filtered out. Normalization method used in DESeq2 was Median Ratio Normalization (MRN) (Maza et al., 2013). The Pearson Correlation Coefficient analysis was performed by ggpubr R package. The Gene Ontology (GO) analysis was performed with DAVID Bioinformatics Resources (v6.8) (Huang da et al., 2009a; b).

cDNA was generated from total RNA using SuperScript III First-Strand Synthesis System (Thermo Fisher Scientific, Cat# 18080051). qPCR was performed with biological and technical triplicates using PowerUp SYBR Green Master Mix (Thermo Fisher Scientific, Cat# A25742) on QuantStudio 3 Real-Time PCR System. Gene expression was normalized to ribosomal protein *rp49*. P values were calculated using the values of ΔCq (Cq (gene of interest) - Cq(rp49)) (Yuan et al., 2006). The sequences of the primers are listed in the Key Resources Table.

### DNA binding analysis and ChIP-qPCR

Overlap in the DNA binding locus for Yki, Bon, and Sd was analyzed and visualized by ChIPpeakAnno using published ChIP-seq datasets of Yki (GSE38594) and Bon (GSE25921) after converting to dm6 coordinates, and DamID-seq dataset of Sd (GSE120731) (Negre *et al*., 2011; Oh *et al*., 2013; Vissers *et al*., 2018; Zhu et al., 2010). The p values between every two datasets were determined by the hypergeometric test using totalTest number for number of potential peaks (Zhu *et al*., 2010). The totalTest number here (89144) was calculated by dividing the *Drosophila* genome (143.7 Mb) (dos Santos et al., 2015) with the largest mean peak width from three datasets (1612 bp for Sd DamID), assuming that these three proteins can potentially bind anywhere in the genome. The presence of ChIP peak and DamID peak at Bon-Yki jointly regulated genes was analyzed using IGV (Robinson et al., 2011).

ChIP assay was performed using Chromatin Immunoprecipitation (ChIP) Assay Kit (MilliporeSigma, Cat# 17-295) with adaptations based on the application notes on GFP- and RFP-Trap by ChromoTek. 100 Eye-antennal discs for each replicate were dissected in ice-cold 1xPBS. Protein and DNA were crosslinked by incubating the discs in 1.8% (v/v) formaldehyde (Thermo Fisher Scientific, Cat# 28906) in 1x PBS for 15 min at RT and quenched with 225 mM Glycine for 5 min at RT (Oh *et al*., 2013). Discs were washed three time with 1x PBS, transferred to 200 μl SDS lysis buffer (per 100 discs) containing 2.5 x EDTA-free mini cOmplete Protease Inhibitor (MilliporeSigma, Cat# 11836170001, 1 tablet per 4 ml SDS lysis buffer), homogenized with Pellet Pestle for 30 secs on ice, and incubated for 10 min on ice. The lysate was sonicated using Bioruptor sonicator for 15 min (30 secs on and 30 secs rest at high power) in 4°C water bath to shear the DNA. Samples were centrifuged at 16,000 rcf for 10 min at 4°C, and the supernatant was diluted 10-fold in ChIP Dilution Buffer containing 2.5 x mini cOmplete inhibitor. Samples were pre-cleared with 75 μl (50% slurry) of Pierce Control Agarose Resin for 1 hr at 4°C with rotation, and after a brief centrifugation, 5% sample was saved as input sample. The supernatant was then incubated with 25 μl (50% slurry) of RFP-Trap agarose overnight at 4°C with rotation. The agarose beads were washed as indicated in the kit. The protein-DNA complex was eluted twice with 250 μl elution buffer for 15 min, first at RT on a nutator and second on a 65°C shaker. Input samples and elutes were reverse cross-linked with 0.2 M NaCl at 65°C overnight, digested with RNase A (Thermo Fisher Scientific, Cat# EN0531) for 30 min at 37°C and Proteinase K (Thermo Fisher Scientific, Cat# 26160) as indicated in the kit. DNA was recovered by phenol/chloroform extraction and ethanol precipitation.

qPCR was performed with biological triplicates and technical duplicates using PowerUp SYBR Green Master Mix on QuantStudio 3 Real-Time PCR System. Ratio to the input was presented. P values were calculated using the values of ΔCq (Cq (ChIPped) - Cq(input)). The sequences of the primers are listed in the Key Resources Table.

### Quantification And Statistical Analysis

Statistical significance of categorical variables was determined by Fisher’s exact test with two-sided p value calculation using GraphPad Prism (v9.1.0). Unless indicated otherwise, in all other statistical analyses, data were presented as mean ± SD of at least three replicates, and the significance was determined by unpaired t-test with Welch’s correction using GraphPad Prism (v9.1.0). The numbers (n) and values of each sample used per experiment are provided in Table S5. The p values are presented as follows: ns (p > 0.05, not significant), * (p ≤ 0.05), ** (p ≤ 0.01), *** (p ≤ 0.001), **** (p ≤ 0.0001).

## Supplemental Tables

**Table S1**. Yki interactome (related to Figure 1).

**Table S2**. Bon interactome (related to Figure 5).

**Table S3**. Differential gene expression analysis (related to Figure 6 and S6).

**Table S4**. GO_BP analysis of joint targets of Bon and Yki (related to Figure 6).

**Table S5**. Quantifications of phenotypes (related to Figure 2, 3, 4, 5, 6, 7, S1, S2, S3, S4, S5, and S6).

