## Supplemental Figures and Legends for "Hippo pathway and Bonus control developmental cell fate decisions in the *Drosophila* eye"

### Figure S1

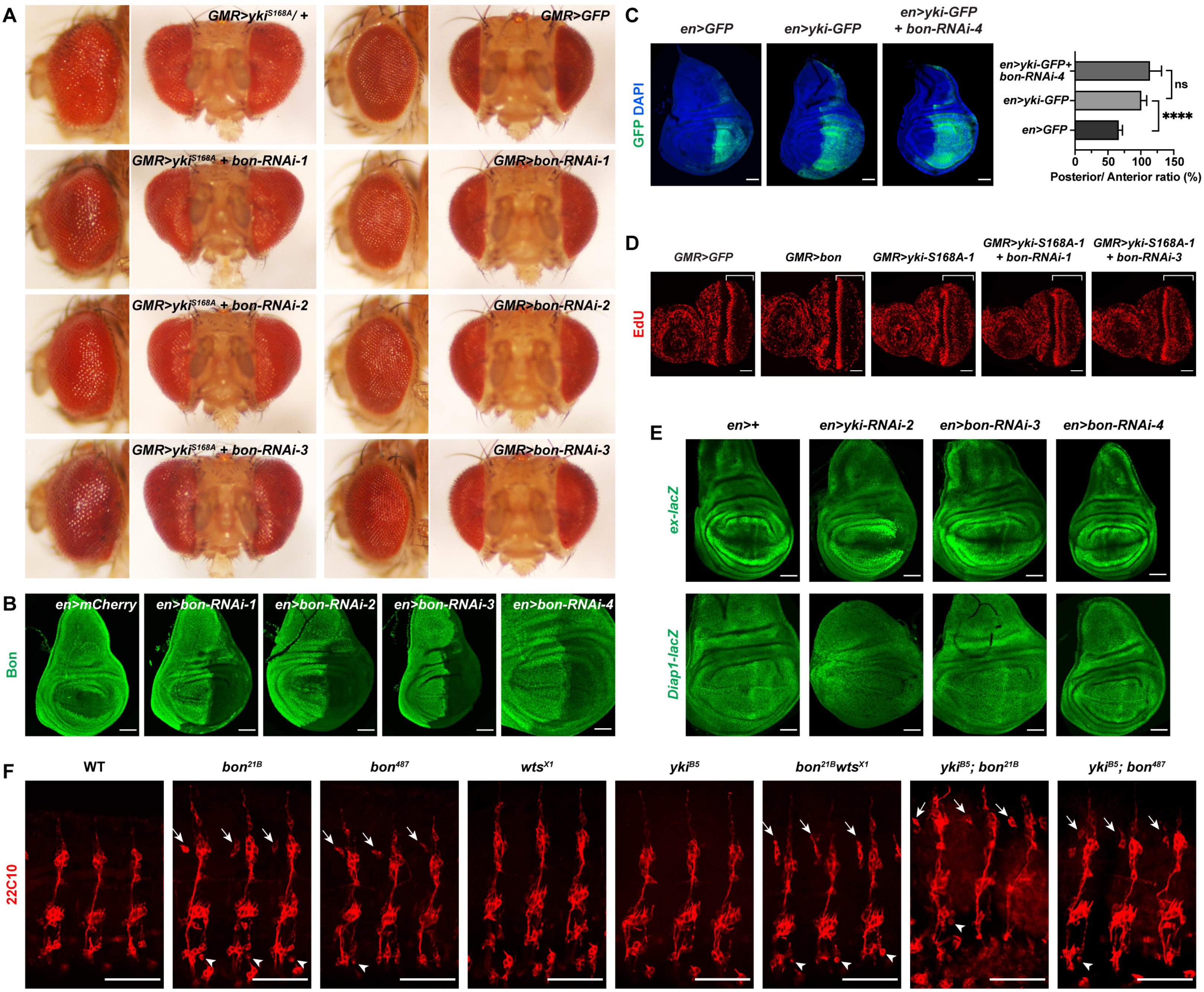
A
GMR>yki S168A/ +
GMR>GFP
GMR>yki S168A + bon-RNAi-1
GMR>bon-RNAi-1
GMR>yki S168A + bon-RNAi-2
GMR>bon-RNAi-2
GMR>yki S168A + bon-RNAi-3
GMR>bon-RNAi-3
B
Bon
en>mCherry
en>bon-RNAi-1
en>bon-RNAi-2
en>bon-RNAi-3
en>bon-RNAi-4
C
en>GFP
en>yki-GFP
en>yki-GFP + bon-RNAi-4
GFP DAPI
en>yki-GFP+ bon-RNAi-4
en>yki-GFP
en>GFP
ns
****
0
50
100
150
Posterior/ Anterior ratio (%)
D
GMR>GFP
GMR>bon
GMR>yki-S168A-1
GMR>yki-S168A-1 + bon-RNAi-1
GMR>yki-S168A-1 + bon-RNAi-3
EdU
E
en>+
en>yki-RNAi-2
en>bon-RNAi-3
en>bon-RNAi-4
ex-lacZ
Diap1-lacZ
F
WT
bon 21B
bon 487
wts X1
yki B5
bon 21B wts X1
yki B5; bon 21B
yki B5; bon 487
22C10

**Figure S1. Independent functions of Yki and Bon in growth regulation and embryonic peripheral nervous system (PNS) development (related to Figure 2).**

(A) Adult eyes expressing indicated *UAS* transgenes with *GMR-GAL4* were examined for growth phenotypes. Knockdown of *bon* with different RNAi lines did not modify Yki-S168A-induced eye overgrowth (left panels) and did not affect eye size when expressed alone (right panels).

(B) Third instar larval (L3) wing discs expressing indicated *UAS* transgenes with the posterior compartment driver *en-GAL4* were immunostained with anti-Bon antibody, confirming the effectiveness of all four Bon RNAi lines. Anterior is on the left. Scale bars: 50 μm.

(C) L3 wing discs expressing indicated *UAS* transgenes with *en-GAL4* were examined for the effect of *bon-RNAi* on Yki-induced overgrowth. Quantification was based on the ratio of the area of the posterior compartment (GFP) to the area of the anterior compartment (non-GFP) of the wing discs. Details of quantification are provided in Table S5. Scale bars: 50 μm.

(D) L3 eye-antennal discs expressing indicated *UAS* transgenes with *GMR-GAL4* were examined for cell proliferation using EdU incorporation assay. Brackets: from the second mitotic wave (SMW) to the posterior end. The ectopic DNA synthesis posterior to the SMW with Yki-S168A overexpression was not affected by knockdown of *bon*. Scale bars: 50 μm.

(E) L3 wing discs expressing indicated *UAS* transgenes with *en-GAL4* were examined for the effect of *bon* RNAi on the reporters of canonical Yki targets, *ex-lacZ* and *Diap1-lacZ*. Scale bars: 50 μm.

(F) Stage 16 embryos of the indicated genotypes were immunostained with neuronal antibody 22C10 to examine the PNS. Oregon R was used as wild-type (WT) control. All mutants were homozygous. Second to fourth abdominal segments (A2-A4) are shown in all images, with anterior to the left and dorsal up. Arrows and arrowheads indicate the ectopic neurons anterior to the dorsal clusters and posterior to the v' clusters, respectively. Scale bars: 50 μm.

### Figure S2

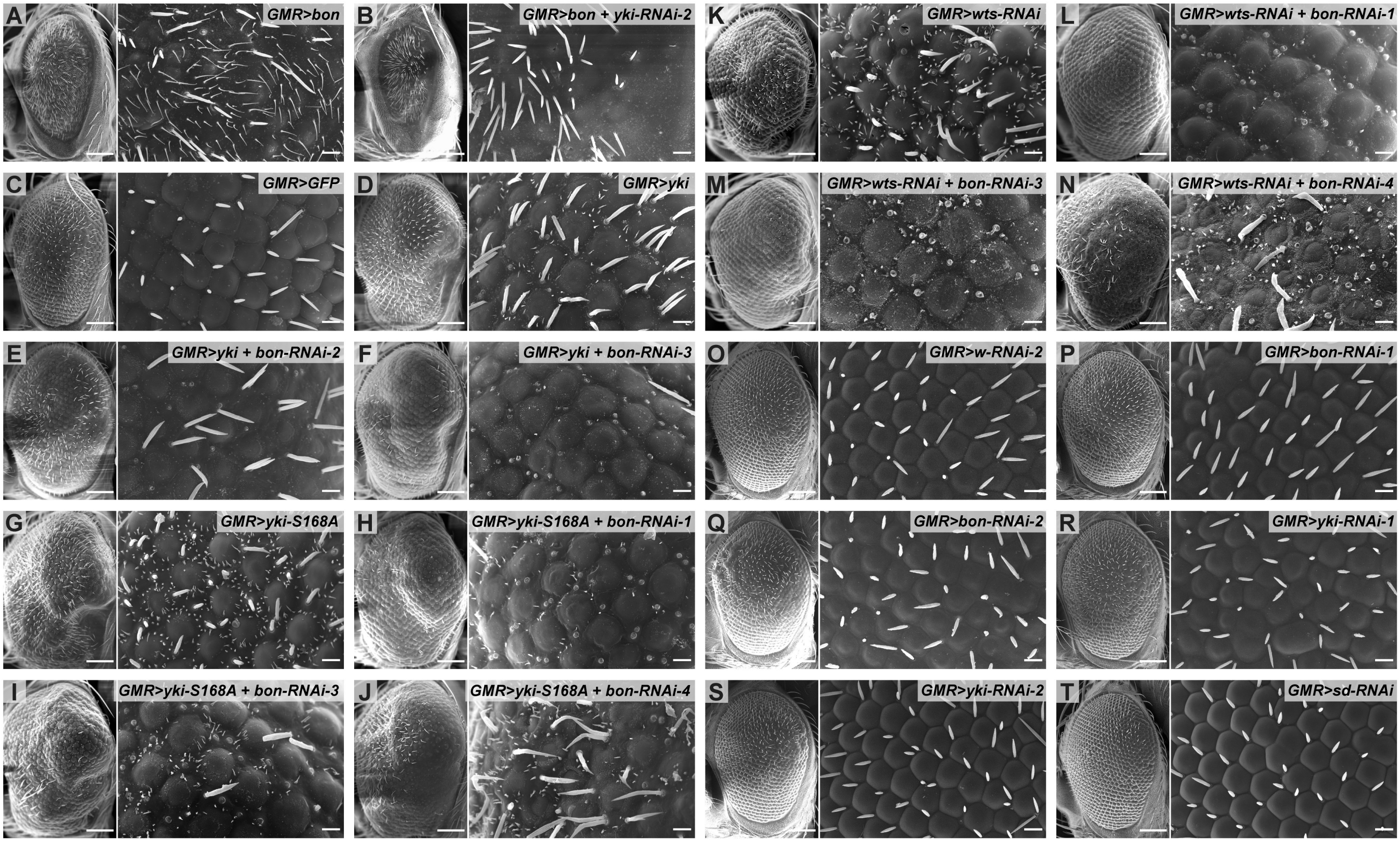
A
GMR>bon
B
GMR>bon + yki-RNAi-2
K
GMR>wts-RNAi
L
GMR>wts-RNAi + bon-RNAi-1
C
GMR>GFP
D
GMR>yki
M
GMR>wts-RNAi + bon-RNAi-3
N
GMR>wts-RNAi + bon-RNAi-4
E
GMR>yki + bon-RNAi-2
F
GMR>yki + bon-RNAi-3
O
GMR>w-RNAi-2
P
GMR>bon-RNAi-1
G
GMR>yki-S168A
H
GMR>yki-S168A + bon-RNAi-1
Q
GMR>bon-RNAi-2
R
GMR>yki-RNAi-1
I
GMR>yki-S168A + bon-RNAi-3
J
GMR>yki-S168A + bon-RNAi-4
S
GMR>yki-RNAi-2
T
GMR>sd-RNAi

**Figure S2. Bon and Yki are mutually dependent for trichome formation in adult eyes (related to Figure 2).**

(A-B) Scanning Electron Microscope (SEM) images of adult eyes with the indicated genotypes, quantified in Figure 2H. Scales bars in left panels: 100 μm; in enlarged views in right panels: 10 μm.

(C-N) SEM images of adult eyes with the indicated genotypes, quantified in Figures 2O-2R or Table S5, showing that *bon-RNAi*s significantly suppressed wild-type Yki (C-F), Yki-S168A (G-J), and *wts* knockdown (K-N) induced trichomes in adult eyes. Crosses were set up at 25°C and shifted to 29°C after the emergence of first instar larvae. Scales bars in left panels: 100 μm; in enlarged views in right panels: 10 μm.

(O-T) SEM images of adult eyes with individual knockdown of *bon*, *yki*, or *sd* with *GMR-GAL4*. *w-RNAi-2* was used as a control. Scales bars in left panels: 100 μm; in enlarged views in right panels: 10 μm.

### Figure S3

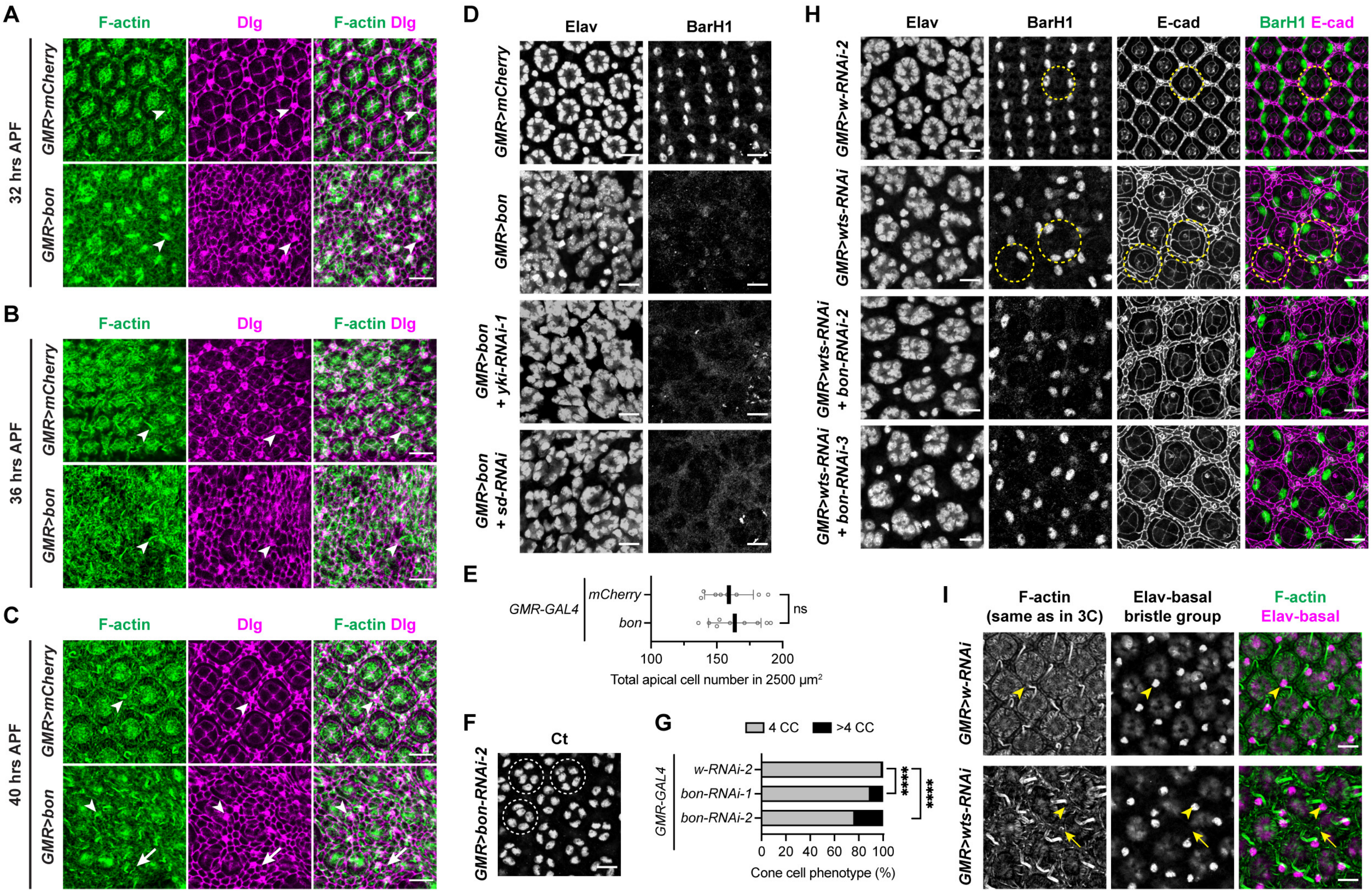

A
B
C
F-actin
Dlg
F-actin Dlg
32 hrs APF
36 hrs APF
40 hrs APF
GMR>mCherry
GMR>bon
D
Elav
BarH1
GMR>bon + yki-RNAi-1
GMR>bon + sd-RNAi
E
GMR-GAL4
mCherry
bon
ns
100
150
200
Total apical cell number in 2500 μm²
F
Ct
GMR>bon-RNAi-2
G
4 CC
>4 CC
w-RNAi-2
bon-RNAi-1
bon-RNAi-2
****
0
20
40
60
80
100
Cone cell phenotype (%)
H
E-cad
BarH1 E-cad
GMR>w-RNAi-2
GMR>wts-RNAi
GMR>wts-RNAi + bon-RNAi-2
GMR>wts-RNAi + bon-RNAi-3
I
F-actin (same as in 3C)
Elav-basal bristle group
F-actin Elav-basal
GMR>w-RNAi
GMR>wts-RNAi

**Figure S3. Bon and Yki promote epidermal trichomes at the expense of retinal cells (related to Figure 3).**

(A-C) Bon induced trichomes are initiated at 40 hrs after puparium formation (APF). Pupal eyes expressing the indicated *UAS* transgenes with *GMR-GAL4* were collected at the indicated time points and stained with phalloidin for F-actin and anti-disc large antibody (Dlg) for cell boundaries. Arrowheads: interommatidial bristles and sockets; arrows: trichomes and the corresponding cells. Scale bars: 10 μm.

(D) Pupal eyes at 44 hrs APF expressing the indicated *UAS* transgenes with *GMR-GAL4* were immunostained with anti-Elav antibody for photoreceptors and bristle groups, or anti-BarH1 antibody for primary pigment cells. Scale bars: 10 μm.

(E) Quantification of the total cell numbers in an apical area of 2500 $\mu m^2$ for 44 hrs APF pupal eyes of the indicated genotypes shown in Figure 3A. All cells outlined by the anti-Dlg staining at the apical surface were counted (photoreceptor cells were not clearly outlined thus not included), including all the cone cells, primary pigment cells, secondary pigment cells, tertiary pigment cells, undifferentiated cells, and bristle groups which were counted as one cell each. Detailed numbers are provided in Table S5.

(F) Pupal eyes at 44 hrs APF with *bon* knocked down using *GMR-GAL4 were* immunostained with anti-Cut (Ct) antibody for cone cells. Dashed circles: individual cone cell cluster per ommatidium. Scale bars: 10 μm.

(G) Quantification of the cone cell numbers per ommatidium for the indicated genotypes, including the one shown in F. The p values were determined using Fisher's exact test between ommatidia with 4 cone cells (CC) per ommatidium and those with >4 CC per ommatidium within indicated comparisons. Detailed numbers are provided in Table S5.

(H) Pupal eyes at 40 hrs APF grown at 29°C (equivalent to 48 hrs APF at 25°C) expressing the indicated *UAS* transgenes with *GMR-GAL4* were immunostained with anti-Elav antibody for photoreceptors and bristle groups, or co-staining with anti-BarH1 and anti-E-cad antibodies for primary pigment cells and cell boundaries, respectively. Dashed circles: individual primary pigment cell clusters or the corresponding ommatidia. Scale bars: 10 μm.

(I) Images of pupal eyes with the indicated genotypes (from Figure 3C) co-stained for F-actin and Elav were analyzed for the difference between trichomes and interommatidial bristles. Elav is expressed in the interommatidial bristle groups (arrowheads) but not in trichomes (arrows). Scale bars: 10 μm.

### Figure S4

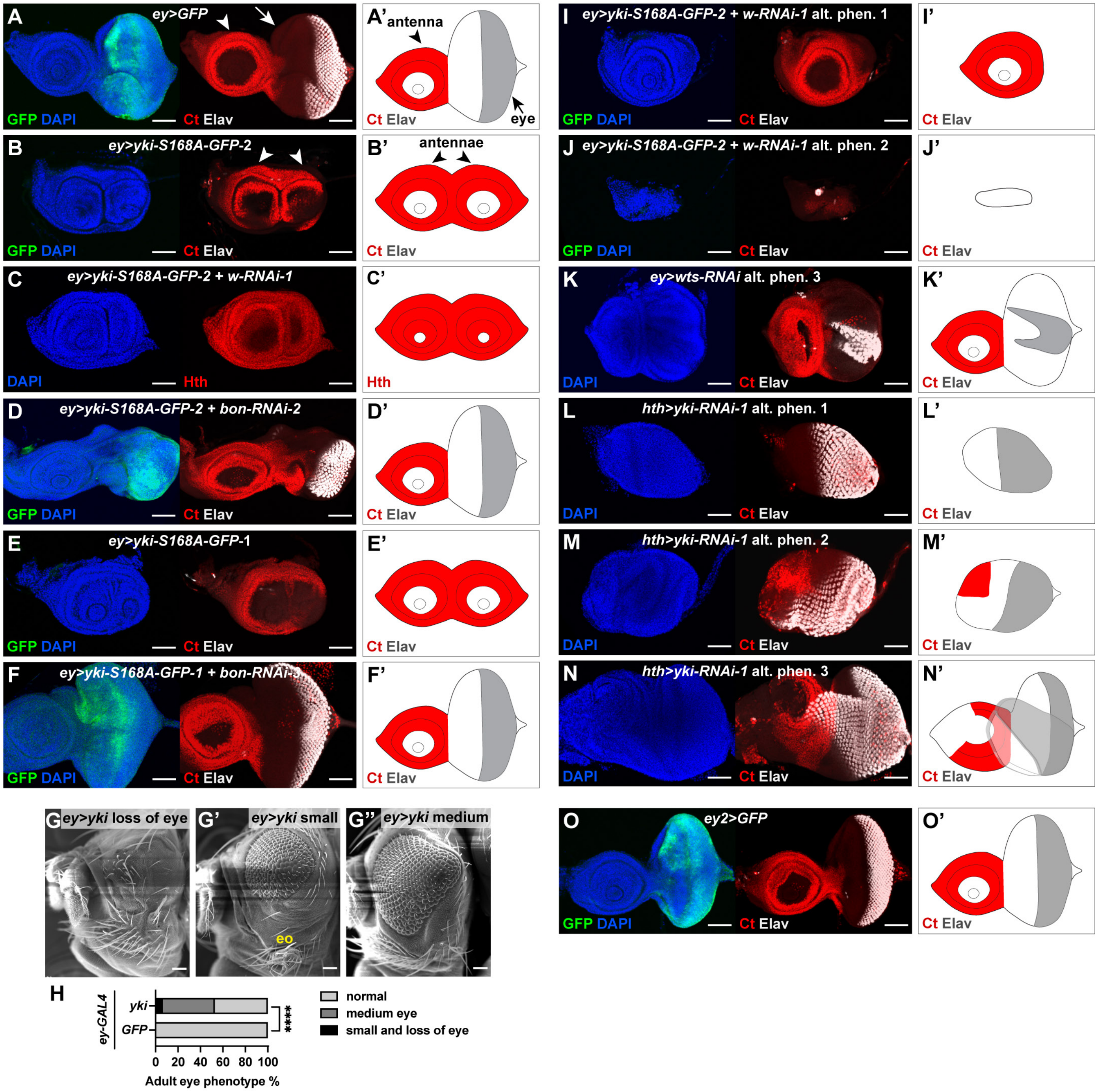

A ey>GFP
GFP DAPI
Ct Elav
A' antenna
eye
B ey>yki-S168A-GFP-2
B' antennae
C ey>yki-S168A-GFP-2 + w-RNAi-1
DAPI
Hth
C'
D ey>yki-S168A-GFP-2 + bon-RNAi-2
D'
E ey>yki-S168A-GFP-1
E'
F ey>yki-S168A-GFP-1 + bon-RNAi-3
F'
G ey>yki loss of eye
G' ey>yki small
eo
G'' ey>yki medium
H
ey-GAL4
yki
GFP
****
0 20 40 60 80 100
Adult eye phenotype %
normal
medium eye
small and loss of eye
I ey>yki-S168A-GFP-2 + w-RNAi-1 alt. phen. 1
I'
J ey>yki-S168A-GFP-2 + w-RNAi-1 alt. phen. 2
J'
K ey>wts-RNAi alt. phen. 3
K'
L hth>yki-RNAi-1 alt. phen. 1
L'
M hth>yki-RNAi-1 alt. phen. 2
M'
N hth>yki-RNAi-1 alt. phen. 3
N'
O ey2>GFP
O'

**Figure S4. Yki and Bon promote antennal fate and epidermal fate while suppressing eye fate (related to Figure 4).**

(A-F') L3 eye-antennal discs with the indicated genotypes (quantified in Figure 4I) were immunostained with anti-Ct or anti-Hth antibody for the antennal compartment and anti-Elav antibody for the neuronal eye fate. (A'-F') Schematic illustrations for the structure of the eye-antennal disc and the pattern of staining in A-F. Arrows: eye discs, arrowheads: antennal discs. Scale bars: 50 μm.

(G-G'') SEM images of adult eyes expressing wild-type Yki with *ey-GAL4*. Representative images are shown for loss of eye (G), small eye (G') and medium eye (G'') phenotypes. eo, epidermal outgrowth. Scale bars: 50 μm.

(H) Quantification of the phenotypes in G-G''. P value was determined using Fisher's exact test between normal and abnormal eye populations within the indicated comparison. The numbers are provided in Table S5.

(I-K') Representative alternative eye disc phenotypes quantified in Figure 4I. (I'-K') Schematic illustrations of I-K. Scale bars: 50 μm.

(L-N') Representative alternative loss of antenna phenotypes quantified in Figure 4P. (L'-N') Schematic illustrations of L-N. Scale bars: 50 μm.

(O-O') L3 eye-antennal disc expressing GFP with *ey-GAL4*-2 (*ey2*) was immunostained with anti-Ct and anti-Elav antibodies and used as a control for *ey2>bon-mCherry* (Figure 4J). (O') Schematic illustration of O. Scale bars: 50 μm.

### Figure S5

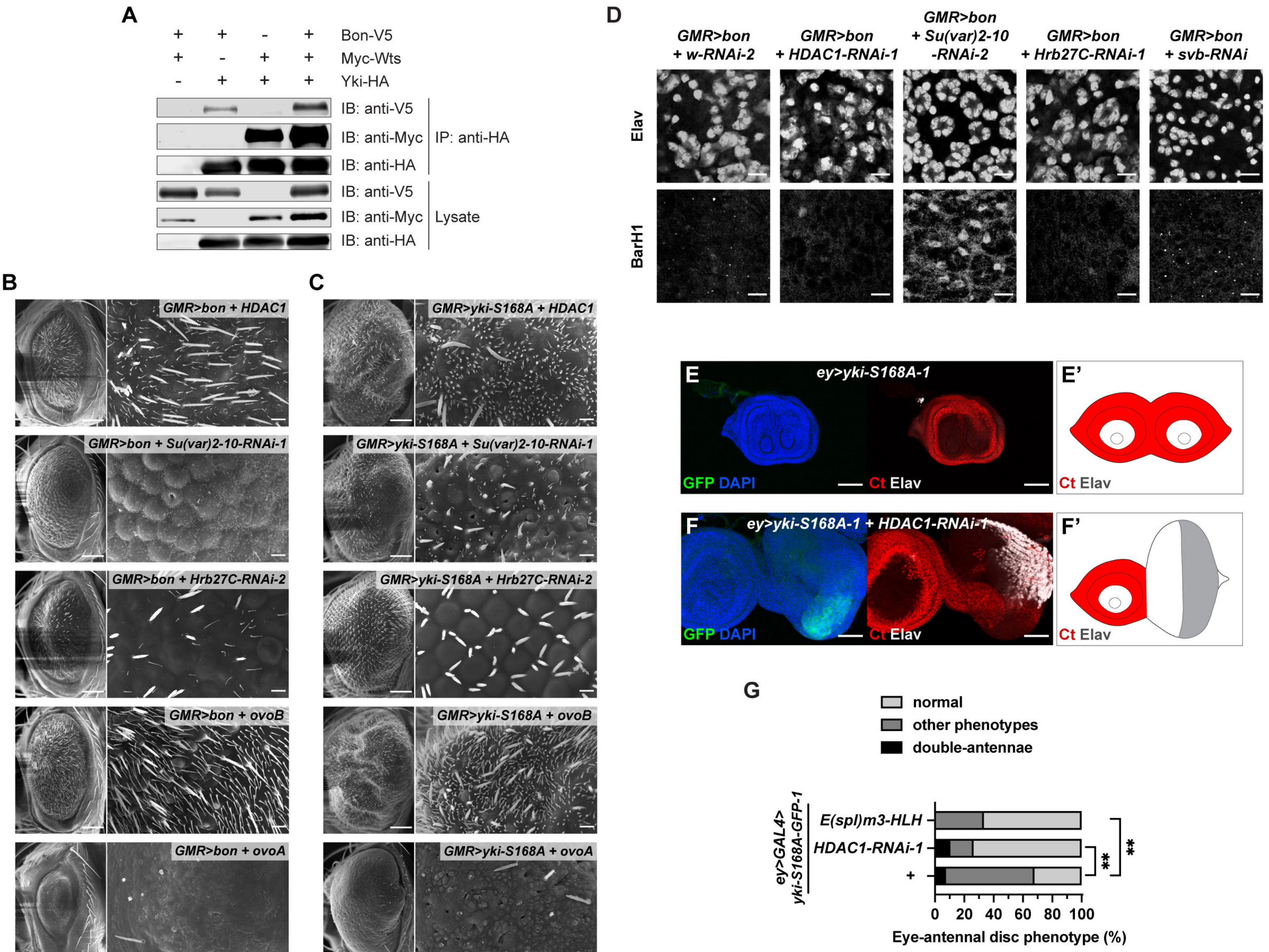
A
Bon-V5
Myc-Wts
Yki-HA
IB: anti-V5
IB: anti-Myc
IB: anti-HA
IP: anti-HA
IB: anti-V5
IB: anti-Myc
IB: anti-HA
Lysate
B
GMR>bon + HDAC1
GMR>bon + Su(var)2-10-RNAi-1
GMR>bon + Hrb27C-RNAi-2
GMR>bon + ovoB
GMR>bon + ovoA
C
GMR>yki-S168A + HDAC1
GMR>yki-S168A + Su(var)2-10-RNAi-1
GMR>yki-S168A + Hrb27C-RNAi-2
GMR>yki-S168A + ovoB
GMR>yki-S168A + ovoA
D
GMR>bon + w-RNAi-2
GMR>bon + HDAC1-RNAi-1
GMR>bon + Su(var)2-10 -RNAi-2
GMR>bon + Hrb27C-RNAi-1
GMR>bon + svb-RNAi
Elav
BarH1
E
ey>yki-S168A-1
GFP DAPI
Ct Elav
E'
Ct Elav
F
ey>yki-S168A-1 + HDAC1-RNAi-1
GFP DAPI
Ct Elav
F'
Ct Elav
G
normal
other phenotypes
double-antennae
ey>GAL4> yki-S168A-GFP-1
E(spl)m3-HLH
HDAC1-RNAi-1
+
**
**
0 20 40 60 80 100
Eye-antennal disc phenotype (%)

**Figure S5. Bon and Yki control eye-epidermal fate determination through cofactors (related to Figure 5).**

(A) Co-IP of Bon-V5, Myc-Wts, and Yki-HA expressed in S2 cells, pulling down with Yki-HA.

(B-C) SEM images of adult eyes with the indicated genotypes (quantified in Figure 5L). Crosses with Bon were kept at 25°C (B), and crosses with Yki-S168A were set up at 25°C and shifted to 29°C after the emergence of first instar larvae (C). Scale bars in left panels: 100 µm; in enlarged views in right panels: 10 µm.

(D) Pupal eyes at 44 hrs APF expressing the indicated *UAS* transgenes with *GMR-GAL4* were immunostained with anti-Elav antibody for photoreceptors and bristle groups, or anti-BarH1 antibody for primary pigment cells. Scale bars: 10 µm.

(E-F') L3 eye-antennal discs expressing the indicated *UAS* transgenes with *ey-GAL4* were immunostained with anti-Ct antibody for the antennal compartment and anti-Elav antibody for the neuronal eye fate. (E'-F') Schematic illustrations of E-F. Scale bars: 50 µm.

(G) Quantification of the phenotypes for genotypes in E-F and Figures 7J-K. P values were determined using Fisher's exact test between normal and abnormal eye disc populations within the indicated comparisons. Detailed numbers are provided in Table S5.

### Figure S6

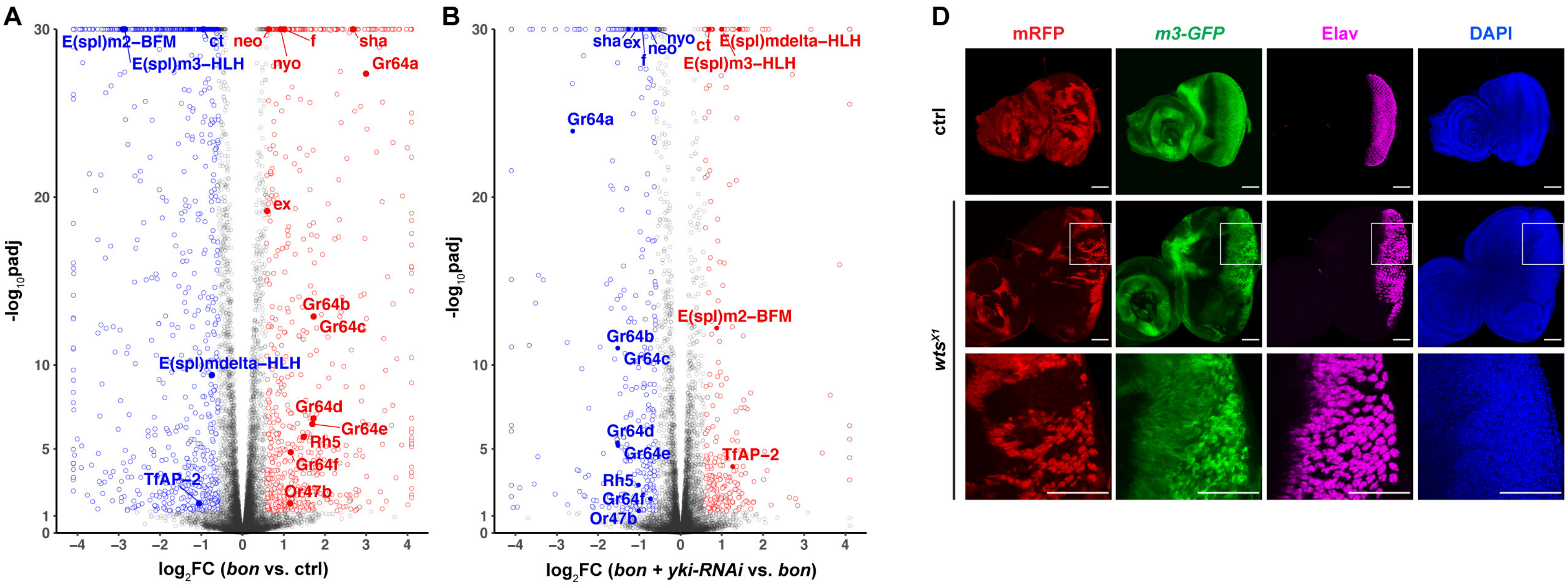


C

| | Yki ChIP peak embryo[1] | Yki ChIP peak wing disc[1] | wts[P2] RNAseq wing disc[1] | Sd DamID peak L3 CNS[2] | Bon ChIP peak embryo[3] |
|---|---|---|---|---|---|
| *sha* | No | No | No | No | No |
| *f* | No | No | Enhanced | No | No |
| *neo* | No | Yes | No | No | No |
| *nyo* | No | No | Enhanced | No | No |
| *Gr64s* | No | No | No | No | No |
| *Or47b* | No | No | No | No | No |
| *E(spl)mdelta-HLH* | Yes | Yes | Repressed | Yes | One peak in this region |
| *E(spl)m2-BFM* | No | Yes | Repressed | Yes | |
| *E(spl)m3-HLH* | Yes | Yes | No | No | |
| *Ct* | Yes | Yes | Repressed | Yes | No |
| *TfAP-2* | Yes | Yes | Repressed | Yes | No |

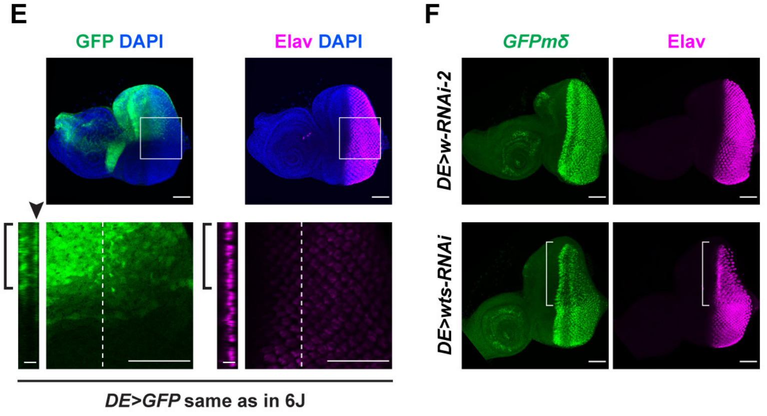


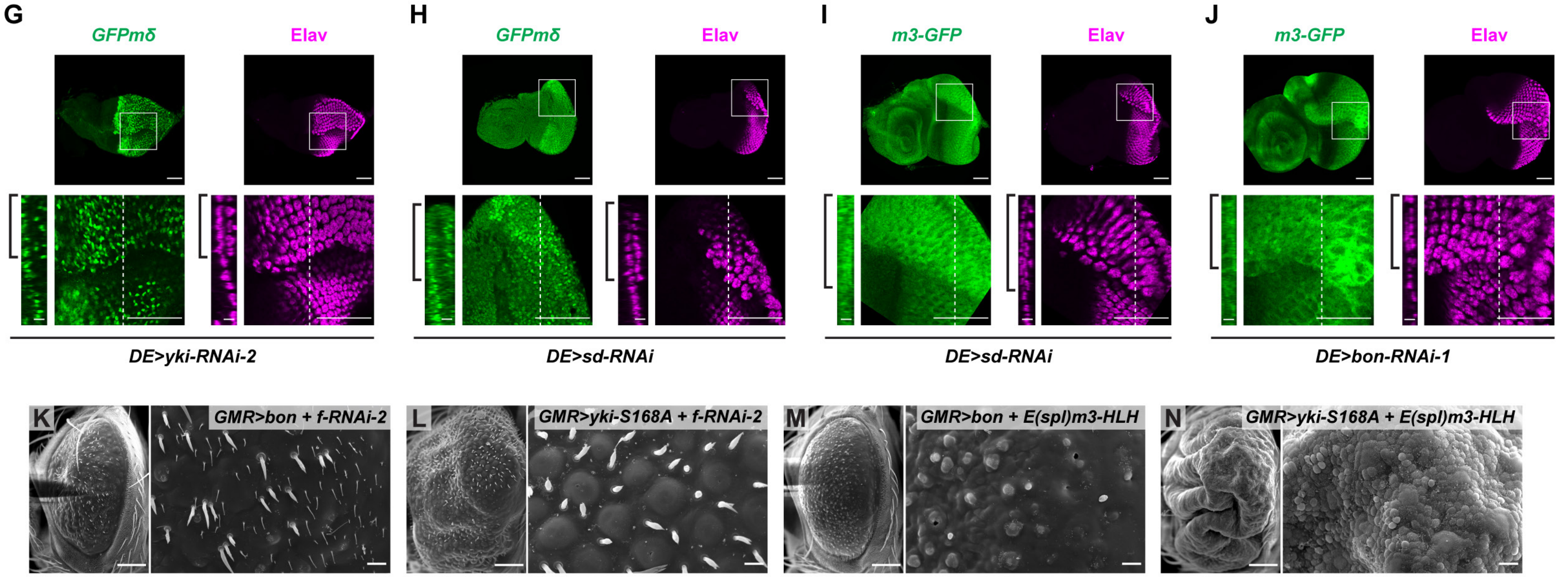

**Figure S6. Bon and Yki control the eye-antenna-epidermis fate determination through their joint transcriptional targets (related to Figures 6 and 7).**

(A-B) Volcano plots showing differentially expressed genes in *GMR>bon-mCherry* compared to *GMR>mCherry* (ctrl) (A) and *GMR>bon-mCherry* + *yki-RNAi-1* compared to *GMR>bon-mCherry* (B) Changes in gene expression were analyzed by DEseq2, and the genes that passed a 1.5 fold change (FC) cutoff and $p$ adj. ≤ 0.05 are highlighted with color. Red circles: significantly upregulated genes. Blue circles: significantly downregulated genes. Solid circles with labels: genes of interest.

(C) Analysis of published ChIP-seq, DamID-seq and RNA-seq datasets for jointly regulated target genes of Bon and Yki. Bon-activated/Yki-dependent genes are in red, and Bon-repressed/Yki-dependent genes are in blue. The assignment of Yes or No for ChIP-seq and DamID-seq was based on the presence of a called peak from 1 kb upstream of the transcription start site to the end of the gene body. The interpretation of the $wts^{P2}$ RNA-seq data was based on the criteria given in the original paper. The following datasets are indicated with superscript numbers: 1. Yki ChIP-seq from 8-16 hrs embryos and L3 wing discs, and RNA-seq from L3 wing discs isolated from the $wts^{P2}$ mutant (GSE38594) (Oh et al., 2013). 2. DamID-seq of Sd from L3 central nervous system (GSE120731) (Vissers et al., 2018). 3. Bon ChIP-seq from 16-24 hrs embryos (GSE25921) (Negre et al., 2011).

(D) L3 eye-antennal discs with control or $wts^{X1}$ mosaic clones were immunostained with anti-GFP antibody for *m3-GFP* reporter and anti-Elav antibody for neuronal eye fate. DAPI was used to ensure that the loss of reporter and Elav was not due to the loss of cells. Mutant clones were generated with *eyFLP* and marked by loss of mRFP. Enlarged views of the boxed regions in $wts^{X1}$ clone are shown in the bottom panels. Scale bars: 50 μm.

(E) L3 eye-antennal disc expressing *UAS-GFP* with *DE-GAL4* (WT control) was immunostained with anti-Elav antibody for neuronal eye fate. Bottom-right panels: enlarged views of the boxed regions with focus set at the peripodial epithelium (PE), bottom-left panels: orthogonal sections at the dashed lines. Brackets: dorsal compartment expressing GFP in both disc proper (DP) and PE. Arrowhead: PE side. The orthogonal sections and their scale bars were scaled 2x along the z axis for easier visualization. Scale bars: 5 μm in orthogonal views and 50 μm in others.

(F) L3 eye-antennal discs expressing indicated *UAS* transgenes with *DE-GAL4* were immunostained with anti-GFP antibody for *GFPmδ* reporter and anti-Elav antibody for neuronal eye fate. Brackets: loss of *GFPmδ* and Elav in the dorsal compartment with *DE>wts-RNAi*. Scale bars: 50 μm.

(G-J) L3 eye-antennal discs expressing indicated transgenes with *DE-GAL4* were immunostained with anti-GFP antibody for *GFPmδ* or *m3-GFP* reporter and anti-Elav antibody for neuronal eye fate. Top and bottom-right panels are focused at the PE. Bottom-right panels: enlarged views of the boxed regions, bottom-left panels: orthogonal sections at the dashed lines, brackets: gain of *GFPmδ*, *m3-GFP* and Elav in the PE layer of the dorsal compartment. The orthogonal sections and their scale bars were scaled 2x along the z axis for easier visualization. Scale bars: 5 μm in orthogonal views and 50 μm in others.

(K-N) SEM images of adult eyes with indicated genotypes (quantified in Figure 7I). Crosses with Bon were kept at 25°C (K and M), and crosses with Yki-S168A were set up at 25°C and shifted to 29°C after the emergence of first instar larvae (L and N). Scale bars in left panels: 100 μm; in enlarged views in right panels: 10 μm.

### Figure S7

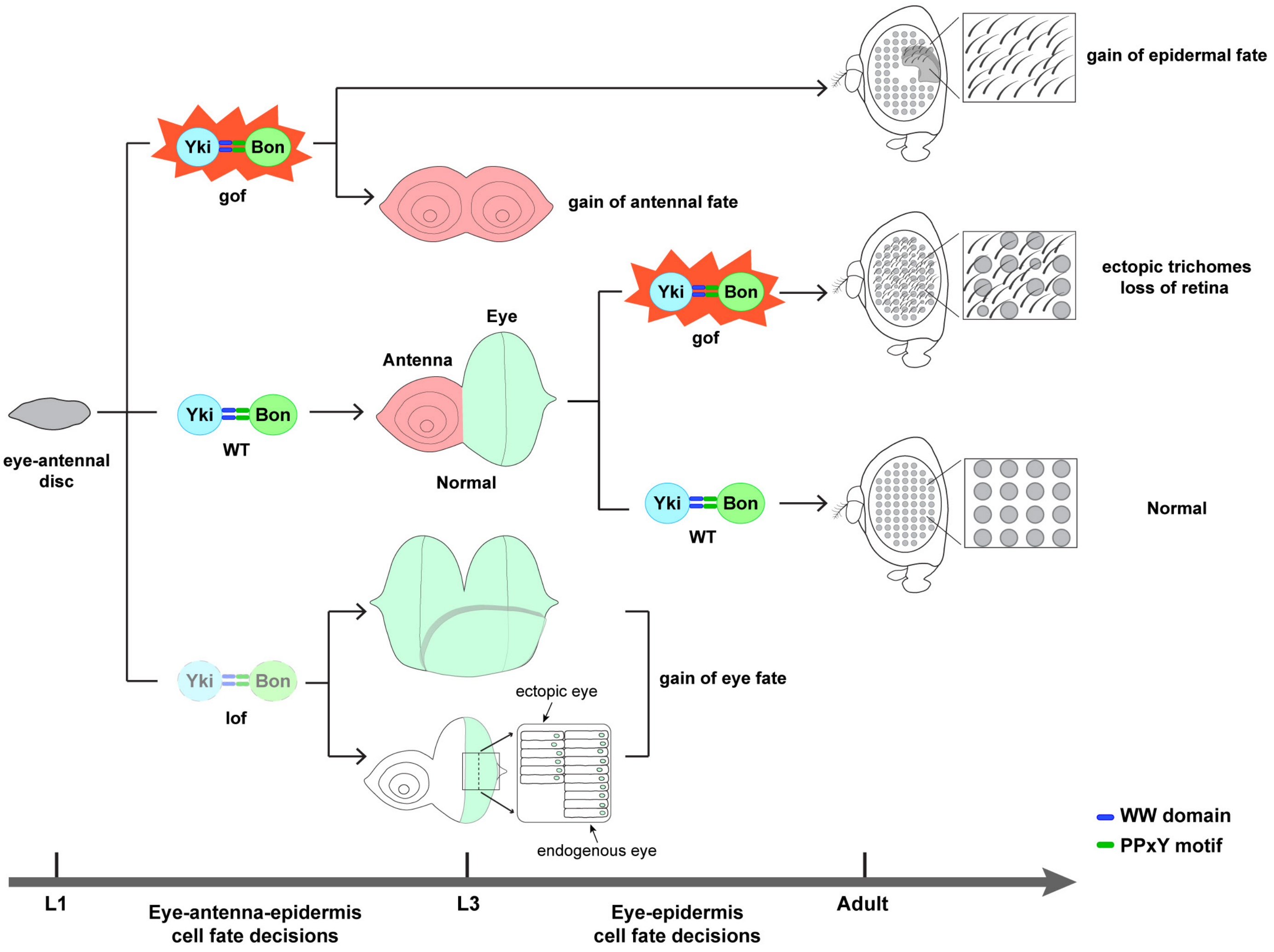

gain of epidermal fate
Yki
Bon
gof
gain of antennal fate
ectopic trichomes
loss of retina
Eye
Antenna
eye-antennal
disc
WT
Normal
Normal
ectopic eye
gain of eye fate
lof
endogenous eye
WW domain
PPxY motif
L1
Eye-antenna-epidermis
cell fate decisions
L3
Eye-epidermis
cell fate decisions
Adult

**Figure S7. The Yki-Bon complex regulates cell fate decisions in the eye at two stages during *Drosophila* eye development (related to Figures 6 and 7).**

First, the Yki-Bon complex promotes antennal and epidermal fates and inhibits the eye fate during the early eye field specification, before the L3 larval stage, whereas Wts counteracts this activity. When the Yki-Bon complex is activated early (gof, gain of function), it results in eye-to-antenna transformation and/or epidermal outgrowth. Early inactivation of the complex (lof, loss of function) results in ectopic eye fate seen as antenna-to-eye transformation or induction of ectopic eye markers in the eye discs at L3. Second, after the segregation of the eye/antenna/epidermis fields and the start of MF in L3, the Yki-Bon complex promotes the epidermal cell fate while suppressing the retinal fate, whereas Wts ensures the proper differentiation of the retina. When the Yki-Bon complex is activated late (gof after L3), it induces ectopic epidermal cells with trichomes at the expense of the retinal cells.
